# Capybara: A computational tool to measure cell identity and fate transitions

**DOI:** 10.1101/2020.02.17.947390

**Authors:** Wenjun Kong, Yuheng C. Fu, Samantha A. Morris

## Abstract

Transitions in cell identity are fundamental to development, reprogramming, and disease. Single-cell technologies enable the dissection of tissue composition on a cell-by-cell basis in complex biological systems. However, highly-sparse single-cell RNA-seq data poses challenges for cell-type identification algorithms based on bulk RNA-seq. Single-cell analytical tools are also limited, where they require prior biological knowledge and typically classify cells in a discrete, categorical manner. Here, we present a computational tool, ‘Capybara,’ designed to measure cell identity as a continuum, at single-cell resolution. This approach enables the classification of discrete cell entities but also identifies cells harboring multiple identities, supporting a metric to quantify cell fate transition dynamics. We benchmark the performance of Capybara against other existing classifiers and demonstrate its efficacy to annotate cells and identify critical transitions within a well-characterized differentiation hierarchy, hematopoiesis. Our application of Capybara to a range of reprogramming strategies reveals previously uncharacterized regional patterning and identifies a putative *in vivo* correlate for an engineered cell type that has, to date, remained undefined. These findings prioritize interventions to increase the efficiency and fidelity of cell engineering strategies, showcasing the utility of Capybara to dissect cell identity and fate transitions. Capybara code and documentation are available at https://github.com/morris-lab/Capybara.

## Introduction

Cells are the fundamental units that form complex biological systems, with each defined cell type assuming a specific function (Altschuler and Wu, 2010). The cataloging of cellular identity is essential for the standardization of cell biology; traditionally, cells have been classified based on features such as their location, morphology, and behavior. More recently, advances in molecular biology have supported more detailed definitions of cell type. Current endeavors to create cell atlases across a range of organisms have reignited interest in the systematic classification of cell identity (Han et al., 2018; Regev et al., 2017; Tabula Muris Consortium et al., 2018). These efforts focus primarily on defining the cell types that comprise adult tissues under homeostasis.

In contrast, cataloging cell identity in dynamic contexts, such as development, disease, and reprogramming, represents a challenge as cell type and state are in continual transition. Indeed, previous computational approaches to assess cell identity based on gene expression measurements (Cahan et al., 2014) suggests that reprogrammed cell identity exists on a continuum, recapitulating only some elements of the target cell type and retaining signatures of the original cell identity (Morris et al., 2014). However, since these measurements were based on bulk expression analysis, the contribution of population heterogeneity to these observations could not be assessed.

Recently emerging high-throughput single-cell RNA-sequencing (scRNA-seq) technologies have revolutionized the resolution of sequencing to a new standard (Klein et al., 2015; Macosko et al., 2015; Zheng et al., 2017). This rapidly evolving technique has opened broad opportunities to assess cell composition and dynamics in complex systems. Accompanying the broad adoption of these new technologies is a demand for methods to automate cell type classification; for recently established cell atlases assembled using single-cell profiling, cells are typically manually annotated via unsupervised clustering and known cell-type-specific marker analysis for each cluster (Han et al., 2018; Tabula Muris Consortium et al., 2018). However, manual annotation can be time-consuming and inaccurate, especially if prior knowledge of cell-specific markers is limited.

Several computational approaches have recently emerged to support the automated annotation of cell identity from single-cell data (Abdelaal et al., 2019). For example, Garnett leverages both scRNA-seq and scATAC-seq data to classify single-cell identity (Pliner et al., 2019). In contrast, ScPred uses scRNA-seq alone to build a prediction model based on a training dataset, estimating the probability of each cell belonging to a cell type category (Alquicira-Hernandez et al., 2019). Furthermore, SingleCellNet is an approach that quantitatively assesses identity using a Random Forest classifier to learn cell type-specific gene pairs from cross-platform and cross-species datasets (Tan and Cahan, 2019). However, many of these current supervised methods require prior biological knowledge. Moreover, these approaches deliver discrete, categorical identification of cell type, which may overlook cells that are in fate transitions or represent mixed identities. Indeed, Tan and Cahan, 2019, note the existence of individual cells that harbor dual identity in their classification results.

While discrete cell cataloging is beneficial in describing cells with defined, stable identities, it is limited for the evaluation of cells in continuous biological processes, such as embryonic development and reprogramming. Here, we present Capybara, an unsupervised method to quantitatively assess cell identity as a continuous property. In contrast to current approaches to annotate cell identity, we designed Capybara to interrogate the gradual transition of cell identity during cell differentiation, reprogramming, and disease progression. To achieve this, we measure the identity scores for each query cell against an exhaustive public cell type collection using quadratic programming, a method previously used to evaluate cell progression between defined cell identities in direct lineage reprogramming (Biddy et al., 2018; Treutlein et al., 2016). This approach scores cell identity on a continuum, as a linear combination of all potential identities. We use these continuous identity scores to determine the discrete cell class of each cell. However, in contrast to other current methods, Capybara allows multiple identities to be assigned to an individual cell, enabling the identification of transitioning cells. We build on this unique feature of Capybara to develop a ‘transition metric,’ enabling the characterization of cell fate transition dynamics for a biological process. The efficacy and robustness of Capybara as a classifier is evaluated based on its ability to assign cell types that match manual annotations and benchmarked using a comprehensive assessment method (Abdelaal et al., 2019).

Here, we demonstrate the efficacy of Capybara to classify cell identity and highlight key transition states in a well-studied developmental process, hematopoiesis. We also apply Capybara to characterize cell identity and fate transitions across a broad range of cell reprogramming paradigms. These analyses reveal the absence of regional patterning in the generation of motor neurons from embryonic stem cells (ESCs), and a previously uncharacterized chamber specification event in cardiac fibroblast to cardiomyocyte reprogramming. Finally, our analysis of direct reprogramming from fibroblast to induced endoderm progenitors (iEPs), defines previously unanticipated cell state transitions and identifies an *in vivo* correlate for this relatively uncharacterized reprogrammed cell type. Together, these results showcase the utility of Capybara to dissect cell fate transitions in development and reprogramming, prioritizing new strategies to enhance the fidelity of cell engineering. Capybara code and documentation are available via https://github.com/morris-lab/Capybara.

## Results

### Overview of the Capybara pipeline

Capybara is rooted in previous approaches designed to measure continuous changes in cell identity during direct lineage reprogramming, using Quadratic Programming (QP; see methods), a method that uses bulk expression signatures to classify single cells as a linear combination of different cell types (Biddy et al., 2018; Treutlein et al., 2016). Under this model, each cell receives a continuous score for defined cell types, enabling cell identity to be quantified. Although this method yields a general picture of the cell-type composition of a single-cell dataset, the requirement for gene signature selection and low cellular resolution of bulk datasets limits its overall resolution. We have previously adapted this approach to incorporate scRNA-seq datasets, using the entire gene set as a reference, permitting higher resolution cell-type classification (Seiler et al., 2019). Here, we generalize this method via systematic reference construction, using both bulk and annotated single-cell atlas datasets, enabling the unsupervised assignment of discrete and continuous cell identity measurements in the following steps (**Figure 1; Methods**):

**Figure 1.**
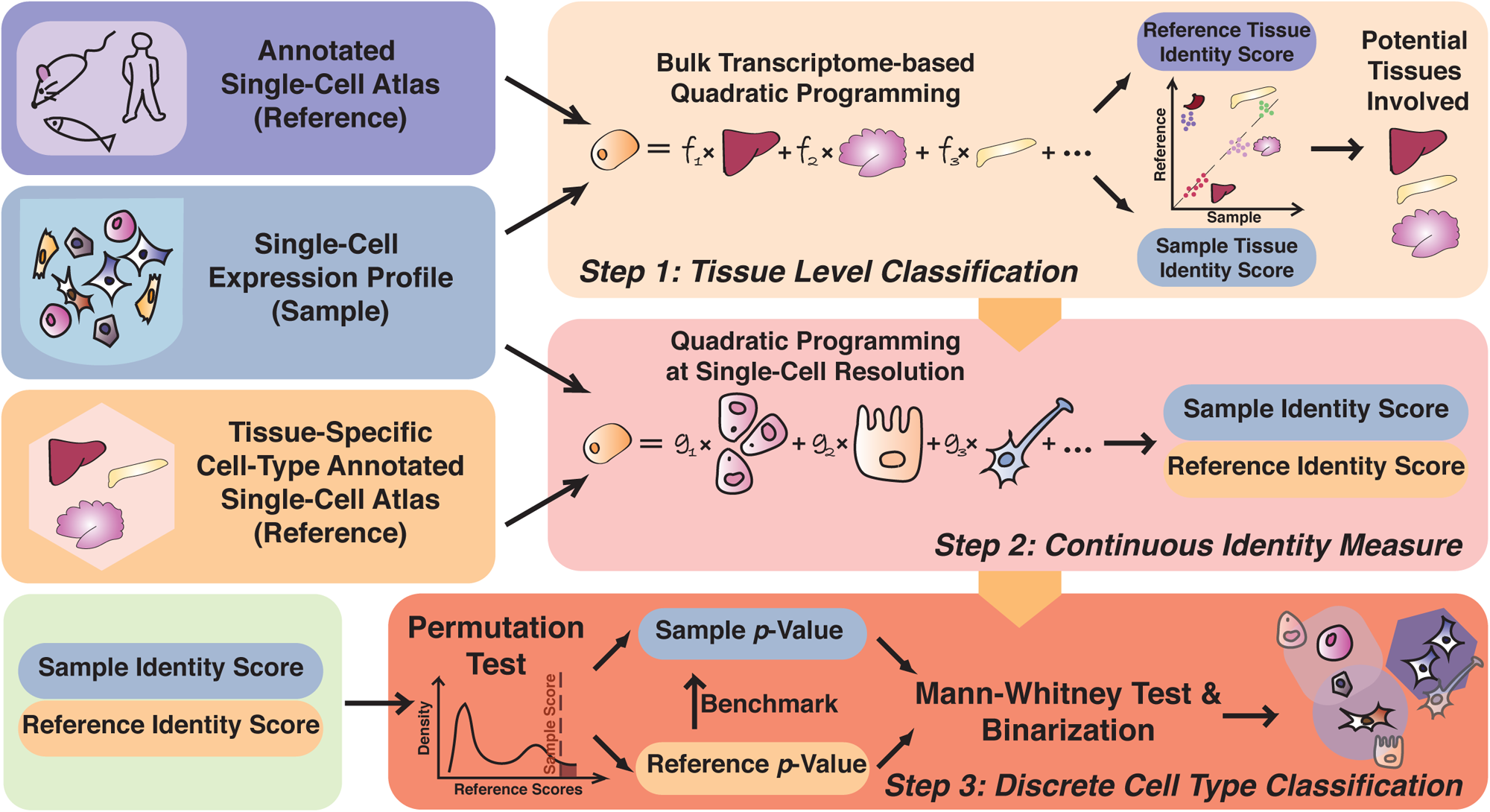
Overview of the Capybara workflow. Left: Input datasets, such as single-cell atlas data and single-cell sample data for evaluation. Right: The three major steps of Capybara pipeline, including tissue-level classification, continuous identity measurement, and discrete cell type classification. In brief, we first perform a tissue-level classification to restrict the number of reference cell types in the downstream analysis. This classification is performed based on bulk transcriptome profiles of different tissues from ARCHS4 (Lachmann et al., 2018). We further identify the correlated tissues in the single-cell atlases, such as the Mouse Cell Atlas (MCA; (Han et al., 2018)). Using only the highly correlated tissues, we build a high-resolution reference. Application of quadratic programming (QP) using this reference provides a continuous measure of cell identity as a linear combination of all cell types within the reference. Finally, we perform discrete classification via randomized testing and score cells with multiple identities using a Mann-Whitney test.

#### Step 1: Tissue-level classification

The performance of Capybara hinges on the selection of an appropriate single-cell reference to classify cell identity. Before assessing cell identity at single-cell resolution, we perform a tissue-level classification designed to restrict the number of reference cell types included in downstream analysis, reducing excessive noise and dependencies caused by correlation across tissues in the final single-cell reference. This tissue-level classification is performed using bulk transcriptomics from ARCHS4, an exhaustive resource platform comprising the majority of published RNA-seq datasets (Lachmann et al., 2018). To construct a relatively comprehensive and clean bulk reference, we filter the available datasets to include only poly-A- or total RNA-seq data from C57BL/6 mice. To further ensure homogeneity in the tissue reference, we use Pearson’s correlation to evaluate the similarity among transcriptome profiles from the same tissue, across different experiments. We then select the top 90 correlated samples to generate the final bulk profile for a total of 30 tissues (**Figure S1A**). Using these tissue transcriptome profiles, QP is applied to the single-cell reference as well as the test sample. Under the assumption that cells from a tissue should receive similar combinations of identity scores, we use a Pearson’s correlation-based metric to align the sample cells to relevant tissues in the single-cell reference. The outcome of this step is the identification of potential tissues present in the sample.

#### Step 2: Generation of a high-resolution custom reference, and continuous identity measurement

Having identified the potential tissues present in a sample from the tissue-level classification, we next assemble a custom single-cell reference dataset, containing the relevant cell types to classify the sample cells accurately. An example of such a reference dataset is the Mouse Cell Atlas (MCA; (Han et al., 2018)), which contains both fetal and adult mouse tissues. For each tissue, it offers a detailed breakdown of its cell types, including the same cell type with different marker genes, offering a high-resolution map of cell type composition. Due to the highly sparse nature of scRNA-seq data, an individual cell transcriptome may not provide a complete representation of a cell type. Therefore, to maintain cellular resolution while increasing transcriptional resolution, we created a pseudo-bulk reference by sampling 90-cells from the population, representing an equal balance of cells with the highest and lowest correlation between each other (see ‘Systematic Reference Construction at Single-Cell Resolution’ in **Methods** for details). This approach creates a pseudo-bulk reference that retains the high-resolution of the single-cell reference, assuming homogeneity in the annotated population of the original single-cell reference. Application of QP using this ‘high-resolution’ reference generates a continuous measurement of cell identity as a linear combination of all cell types within the reference.

#### Step 3: Discrete cell type classification and multiple identity scoring

While continuous identity scores are informative, discrete cell-type assignment offers a more practical assessment of cell-type composition for a biological system. One approach to call discrete cell types is to apply a threshold to the calculated continuous scores. However, threshold selection and quality of the custom high-resolution reference can bias cell type calling via this approach. To overcome this limitation, we apply QP to score cells in the single-cell reference against the bulk reference. This strategy accounts for reference quality, enabling background matrices to be generated, charting the distributions of possible identity scores for each cell type. Then, with identity scores calculated for the sample cells using the high-resolution reference, a randomized test is performed using the background distributions as null to compute the occurrence probability or empirical p-values of each identity score. This test shapes the likelihood identity score occurrence as a continuous distribution, in which the cell type with the lowest likelihood rank is the classified identity. Cross-validations of Capybara using a recent benchmark algorithm (Abdelaal et al., 2019) and our in-house data demonstrate its efficacy to classify discrete cell types accurately (See Method Validation; **Figure S1B, S2**). Overall, using 5 human pancreatic datasets with the benchmark algorithm, the performance of Capybara indicates similar accuracy (rank #5) and median F1 score (rank #5) with reasonable runtime, when benchmarked against 8 other classifiers (**Figure S1B**). This statistical framework also allows us to identify cells that harbor multiple identities, potentially representing cells transitioning between defined cell identities. This function distinguishes Capybara from other current cell-type classifiers, where cells in transition states are classified as either unknown or are assigned definitive identity. To capture multiple cell identities, we use a Mann-Whitney (MW) test to compare the occurrence probabilities of the cell type with the lowest likelihood rank to that of other cell types, following the order from the second-lowest to the highest rank-sum. From this test, we calculate a p-value to determine whether two identities are equally likely to represent the identity of a specific cell. We stop our comparison when we identify the first cell type that is significantly (p-value < 0.05) less likely to represent one of the cell identities. This strategy allows more than one identity to be assigned to a single cell, supporting a broader definition of cell identity, including intermediate cell states in addition to terminal cell identities. This unique aspect of Capybara enables us to explore the establishment and maintenance of cell identity in complex, continuous biological systems.

### Capybara accurately captures cell identity and fate transitions in hematopoiesis

Single-cell transcriptome profiling supports the notion that cell identity exists as a continuum, where cells reside on a complex landscape of many possible states. This concept has been supported by differentiation time-series experiments, capturing snapshots of cell states as they gradually decrease in potential and increase in specialization (Trapnell, 2015). Hematopoiesis provides a well-characterized example of continuous cell state transition. This lineage paradigm serves as a valuable model to understand the general mechanisms of lineage specification (Orkin and Zon, 2008). Recent single-cell profiling presents a more comprehensive picture of heterogeneity in hematopoiesis (Paul et al., 2015; Tusi et al., 2018). From Paul *et al*., 2015, a total of 2,730 myeloid progenitors were enriched from mouse bone marrow and processed via massively parallel single-cell RNA-seq (MARS-seq). Collection at a single time point represents a snapshot of hematopoiesis, capturing distinct transcriptomic states, representing a diverse population of cells (Tusi et al., 2018).

Before classification, we use partition-based graph abstraction (PAGA; (Wolf et al., 2019)) to identify a total of 24 clusters, corresponding to different myeloid populations, which we annotate according to Paul *et al*. (**Figure 2A, B**). Next, we perform an initial tissue-level classification, resulting in a high correspondence of the single-cell data to the bone marrow. We then refer to the Mouse Cell Atlas (MCA) as the single-cell reference dataset in this application, where cell types are manually annotated with further marker-based sub-type annotations (Han et al., 2018). This step deconstructs the initial bone marrow classification into three major tissues: bone marrow, bone marrow (c-Kit), and primary mesenchymal stem cells in the MCA reference (**Figure 2C**). These populations serve as the high-resolution custom reference, supporting continuous identity scoring, followed by discrete classification at the single-cell level to reveal a total of 25 defined cell types across 2430 cells. 0.7% of cells (n = 18) were not assigned an identity and 11.6% of cells (n = 282) received multiple identity scores. To investigate the possibility that these cells with multiple identities arise from cell doublets, we applied DoubletFinder (McGinnis et al., 2019) to this dataset. This analysis suggests that only 4.6% of the multiple identity cell population corresponds to cell doublets (n = 13/282 cells), ruling out doublets as the source of multiple identity signals.

**Figure 2.**
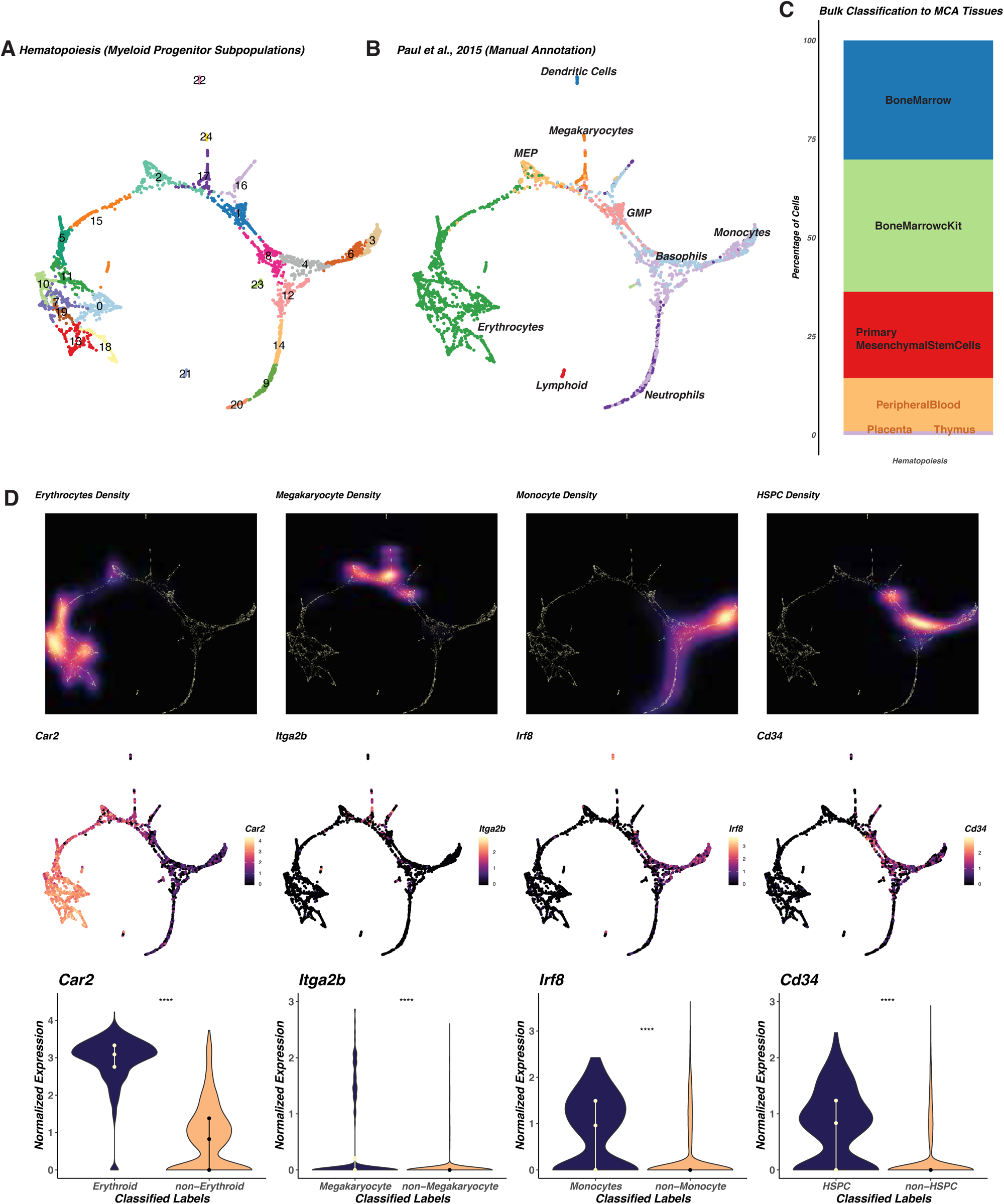
Application of Capybara to classify hematopoietic cell identity. To demonstrate Capybara, we first applied this analysis to a well-characterized paradigm of cell differentiation: hematopoiesis. **(A)** PAGA-guided clustering of an existing hematopoiesis dataset (Paul et al., 2015), consisting of a total of 2,730 myeloid progenitors enriched from mouse bone marrow, revealing a total of 24 clusters. **(B)** Manual annotation of cluster identity, based on Paul et al., 2015. **(C)** In the first step of the Capybara workflow, we perform tissue-level classification of this hematopoiesis dataset using ARCHS4, identifying the correlated tissues in the Mouse Cell Atlas MCA. Major tissues selected in this step include bone marrow, bone marrow (c-Kit), and primary mesenchymal stem cells. **(D)** Capybara classified major populations projected onto the PAGA-guided clustering. Erythrocytes: *Car2*, Megakaryocytes: *Itga2b*, Monocyte: *Irf8*, HSPC: *Cd34*, are used as key markers for different populations. Middle panels: Projection of log-normalized expression of the marker genes. Bottom panels: Comparison of log-normalized expression between classified cell type and other cells. A Wilcoxon test was used for significance testing (****: p <= 0.0001, ***: p <= 0.001, **: p <= 0.01, *: p <= 0.05, ns: p > 0.05).

Overall, Capybara-based cell type classification identified the expected myeloid progenitor populations, including erythrocyte, megakaryocyte, hematopoietic stem/progenitor cells (HSPC), monocytes and neutrophils, agreeing with the original cell annotations (**Figure 2D;** (Paul et al., 2015)). Moreover, expression of known cell type markers defining this hierarchy (*Car2, Itga2b, Cd34*, and *Irf8*) are significantly enriched in their respective cell types (*P* < 2.2E-16), confirming the efficacy of our classification method (**Figure 2D**). Altogether, the application of Capybara to this dataset accurately identifies major hematopoietic cell populations. We next turned our attention to cells with multiple identities to investigate potential transition states within this hierarchy.

### Multiple identity scoring highlights cell fate transitions, supporting a transition metric

Focusing on the ∼12% of cells that harbor multiple identities, projection of these cells onto the PAGA-guided clustering highlights branch points sitting between defined cell identities (**Figure 3A**), suggesting potential fate transitions. We identified 17 major ‘harbors,’ which we define as more than 15 cells sharing the same multiple identities (**Figure 3B**). Two of these harbors are located within the erythroid lineage, and 15 in the myeloid lineage. The most frequently observed transition occurs between two subtypes of erythrocyte progenitors, which marks the transition of *Car2*-expressing erythrocyte progenitors to more differentiated *Hba-a1*-expressing erythrocytes; Arrow 1 in **Figure 3C(I)** marks the harbor sitting between these defined identities. The other observed transition in the erythroid lineage represents megakaryocyte erythrocyte progenitors (MEPs; Arrow 2) that bifurcate to megakaryocyte progenitors and erythrocyte progenitors (**Figure 3C(I)**).

**Figure 3.**
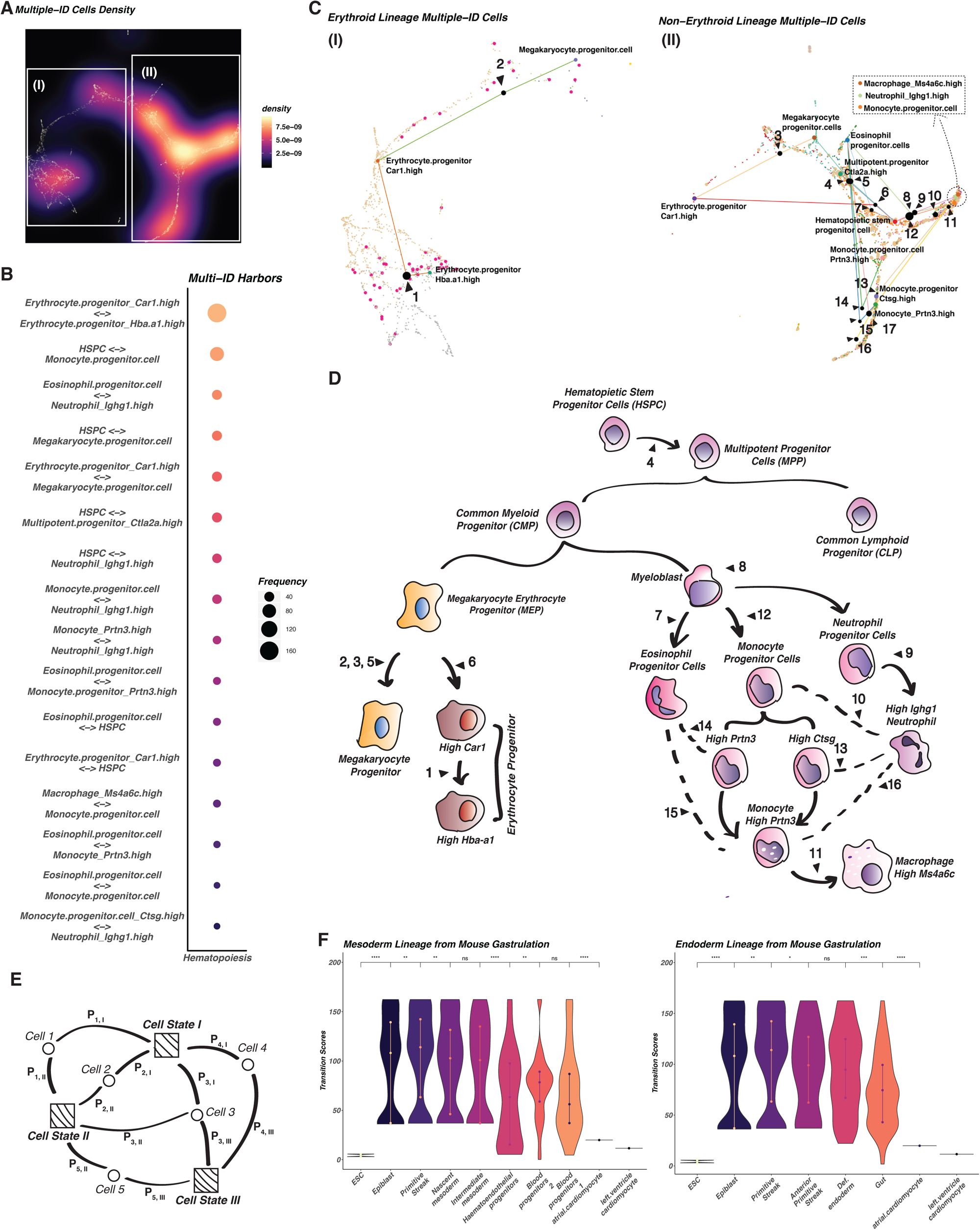
Using Capybara to identify cell fate transitions. A unique feature of Capybara is the quantitative assessment of cell identity as a continuous property, enabling multiple identites to be assigned to an individual cell. We define group of more than 15 cells sharing the same set of multiple identities a ‘harbor’, highlighting branch points sitting between distinct cell identities, suggesting potential fate transitions. **(A)** Density projection of cells with multiple identities onto the PAGA-guided clustering of hematopoiesis (**Figure 2A**), where a lighter color indicates a higher cell density. **(B)** Percentage composition for each combination of multiple identities detected. The size of the dot indicates the percentage of cells with the corresponding set of multiple identities based on the entire population of multiple identity cells. **(C)** Projection of transition harbors onto the PAGA-guided clustering of hematopoiesis. This is the magnified region from **(A)**. The black dots indicate the harbors of the same multiple identity cells, where the size of the dot is proportional to the number of cells contained within each harbor. The erythroid lineage is detailed in **(I)**, outlined in **(A-I)**; the granulocyte/macrophage lineage is featured in **(II)**, as outlined in **(A-II). (D)** Placement of the identified harbors within the hematopoiesis lineage. **(E)** Schematic of the principles underlying Capybara’s transition metric. Squares represent discrete cell identities/states. Circles represent cells. *P*_*ij*_ represents the probability of cell *i* transitioning to cell type/state *j*. Then, the transition score of each cell type/state is calculated as the accumulated information received from each cell connection. **(F)** Demonstration of transition scores using the mouse gastrulation atlas (Pijuan-Sala et al., 2019), embryonic stem cell (ESC) dataset (Briggs et al., 2017), and cardiomyocyte dataset (Stone et al., 2019). A Wilcoxon test was used for significance testing (****: p <= 0.0001, ***: p <= 0.001, **: p <= 0.01, *: p <= 0.05, ns: p > 0.05).

In contrast to the erythroid lineage, transitions within the myeloid lineage are more complex, where we observe three types of transition among the 15 harbors. In the first category, we identify known developmental trajectories, including a transition from HSPCs to relatively more differentiated cell states, such as multipotent progenitor cells (MPPs) (Arrow 4) and megakaryocyte progenitors (Arrow 5), and from monocytes to macrophages (Arrow 11) (**Figure 3C(II)**). Additionally, we identify cells transitioning between HSPCs to more differentiated cell types, such as neutrophils, bypassing intermediate progenitor cell types. This observation may be in part due to a lack of appropriate progenitor cell types in the MCA reference. Altogether, employing Capybara to classify hematopoietic cell identity enables us to superimpose intermediate cell states onto the canonical hematopoiesis lineage tree to define transitions in cell identity (**Figure 3D)**. While the majority of these transitions are known, we also note minor transitions between more differentiated states (Arrows 10, 13, 14, 15, 16), such as from eosinophil progenitors to monocytes, invoking previously described lineage switches between closely-related branches (**Figure 3C(II)-D**; (Graf, 2011).

From our application of Capybara to hematopoiesis, the identification of cells harboring multiple identities enables the characterization of transition states. Furthermore, the identities and states that these cells are connected to help define the cell transition milestones within a differentiation hierarchy. Building on this concept, we further leverage the capacity of Capybara to identify cells harboring multiple identities to develop a ‘transition metric.’ Briefly, for each discrete cell identity, we define in a population, we measure the strength and frequency of its connections to mixed identity states (see methods; **Figure 3E**). This generates a metric we define as a ‘transition score,’ where a high score represents a high information state where identities converge, highlighting a putative cell fate transition. This aspect of Capybara is distinct from approaches such as StemID (Grün et al., 2016) and CytoTrace (Gulati et al., 2020) in that our transition score does not aim to quantify potential. Instead, this score aims to identify those cell fate transitions that are central to a developmental/reprogramming process. This approach is also helpful for defining the dynamics of a cell differentiation hierarchy to identify the stages at which a biological system is in flux.

To illustrate the utility of the Capybara transition metric, we measure transition scores across an example developmental time series consisting of several cell types: 1) Embryonic stem cells (ESCs), representing the earliest stages of development. 2) Mouse gastrulation, a pivotal event in development where cells are rapidly specializing (Pijuan-Sala et al., 2019). 3) Cardiomyocytes, representing post-mitotic, terminally differentiated cells. Transition scores for the ESCs are low, as expected; even though these cells have high developmental potential, typical culture conditions minimize their differentiation. In contrast, across gastrulation, transition scores are initially high and decrease as development progresses. We reason that the earlier stages of development involve broad cell fate specification through progenitors higher in the differentiation hierarchy, explaining the initially high score, which decreases as cells subsequently specialize. Finally, terminally differentiated cardiomyocytes receive low transition scores, reflecting their stable identity **(Figure 3F)**. We next apply Capybara to define cell identity and fate transitions in less defined, non-physiological systems, such as cell reprogramming, which aims to manipulate cell identity for disease modeling and therapeutic application.

### Capybara reveals dorsal-ventral patterning deficiencies in motor neuron programming

Different strategies to reprogram cell identity exist: one common approach involves ‘directed differentiation’ of cells from a pluripotent state, in an attempt to recapitulate embryonic development; an alternate strategy aims to directly reprogram cell identity, bypassing developmental states (Cohen and Melton, 2011; Morris and Daley, 2013; Vierbuchen and Wernig, 2011). Previous methods to assess cell identity using bulk classifiers revealed that directed differentiation produced developmentally immature cell types (Cahan et al., 2014). Similarly, direct reprogramming between differentiated states, typically driven by transcription factor (TF) overexpression, was found to yield partially converted cells and off-target identities (Morris et al., 2014). Indeed, incomplete conversion and emergence of unexpected progenitor-like states is a recurring theme across several direct reprogramming strategies (Morris, 2016). However, before the adoption of highly-parallel single-cell profiling, it remained a challenge to tease apart the heterogeneity arising in these cell engineering protocols.

Here, we apply Capybara to assess the identity of engineered cells generated by a diverse range of reprogramming strategies, initially focusing on the generation of spinal motor neurons (MNs) from mouse ESCs. Two distinct methods for MN generation exist: growth-factor-induced differentiation, and TF-induced programming (**Figure 4A**). Induced differentiation involves sequential treatment with Fgfs, Retinoic Acid (RA), and Sonic hedgehog (Shh) (Standard protocol: SP (Wichterle et al., 2002; Wu et al., 2012)), designed to mimic spinal cord development (Sagner and Briscoe, 2019). In TF-mediated direct programming (DP), overexpression of Ngn2, Isl1, and Lhx3 TFs direct ESCs to spinal MNs, a strategy aiming to bypass canonical progenitor states (Mazzoni et al., 2013; Velasco et al., 2017). These two methodologies were recently compared via single-cell profiling, confirming the generation of MNs via these distinctive routes (Briggs et al., 2017).

**Figure 4.**
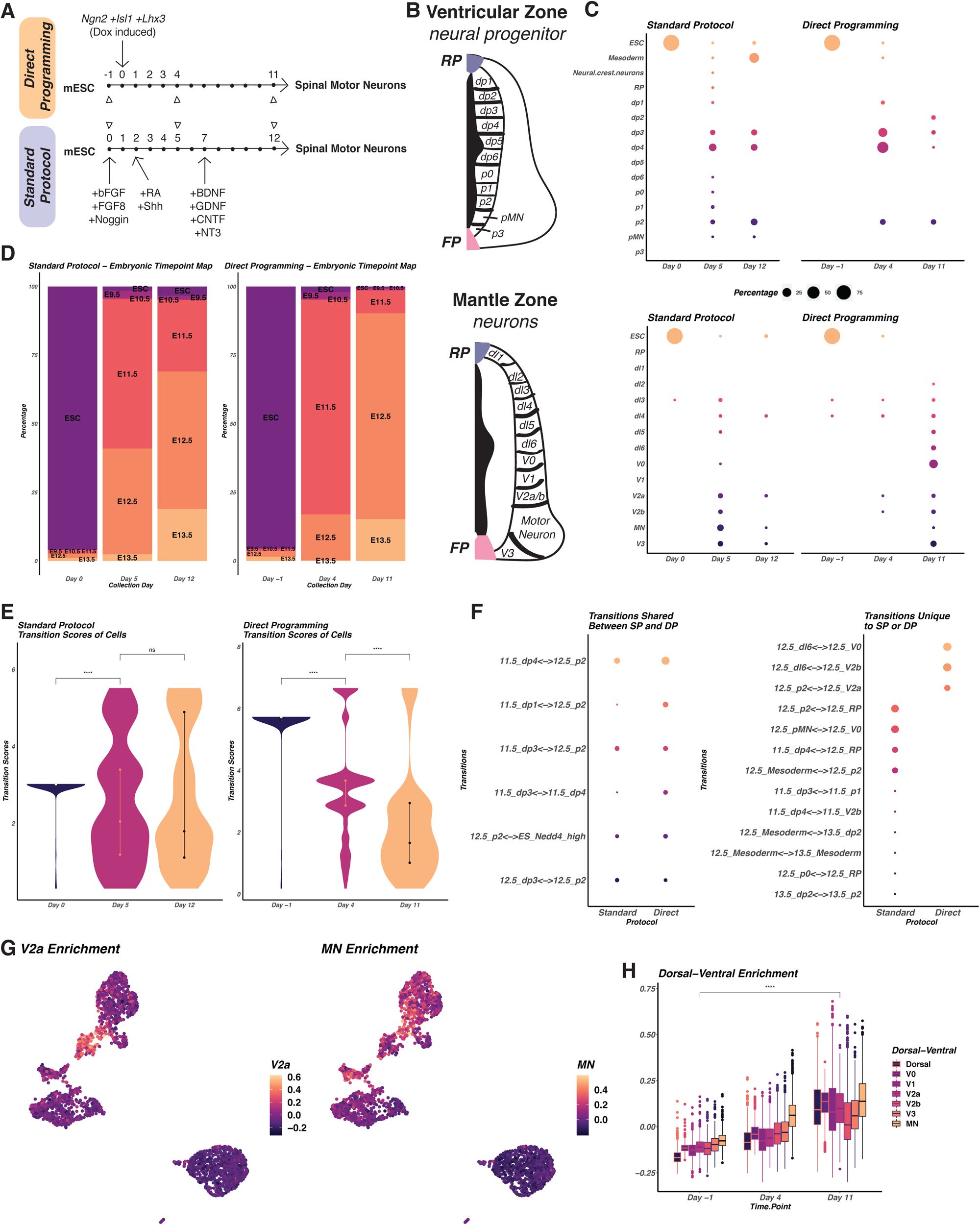
Capybara analysis of spinal motor neuron differentiation and programming. **(A)** Schematic of the experimental protocol, demonstrating differentiation vs. direct programming of motor neurons (MNs) from ESCs, adapted from Briggs et al., 2017. **(B)** Schematic of spinal cord domains and regions. Top: Ventricular zone domains containing neural progenitors. Bottom: Mantle zone domains containing differentiated neurons. **(C)** Capybara analysis showing cell type composition of discrete identities across the two protocols. For each time point, the size of the dot indicates the percent of the overall population contributed by a single cell type. **(D)** Plots depicting the stages of embryonic development that *in vitro*-derived cells map to, divided by protocol and experimental time point. **(E)** Transition scores for each protocol, across experimental time point. A Wilcoxon test was used for significance testing (****: p <= 0.0001, ***: p <= 0.001, **: p <= 0.01, *: p <= 0.05, ns: p > 0.05). **(F)** Transition harbors that are shared between two protocols (left) or are unique to each protocol (right). **(G)** Using known markers (Delile et al., 2019), we calculated enrichment scores for V2a interneuron identity (top) and MN identity (bottom), projecting these scores onto cells from the direct programming protocol, visualized by UMAP. **(H)** Plot of enrichment scores calculated for a range of dorsal-ventral neuron identities, broken down by time point of the direct programming protocol. Significance testing via Wilcoxon testing shows significant increases (p < 2.2E-16) from Day -1 to Day 11 (****: p <= 0.0001, ***: p <= 0.001, **: p <= 0.01, *: p <= 0.05, ns: p > 0.05).

From these single-cell analyses, cells in the standard protocol first commit to the neural lineage via a neural progenitor state, followed by posterior patterning, ventral patterning, and passage through progressive MN progenitor states before the emergence of MNs. Direct programming also involves early neural commitment, but typical spinal embryonic intermediates are bypassed in this scheme, as anticipated. Unexpectedly, the direct approach featured an abnormal intermediate state characterized by a forebrain gene expression signature. Albeit, both protocols successfully generate cells that resemble MNs at the molecular and functional levels (Ichida et al., 2018), with the direct protocol apparently delivering a higher proportion of MNs (Briggs et al., 2017). However, the presence of off-target cell identities, including dorsal spinal cord neurons, muscle, and endoderm, was noted for both approaches, with these unintended cell species dominating the MNs, particularly in the later stages of the standard protocol. Although these detailed characterizations of MN generation from ESCs have revealed valuable information on programming trajectories, the cell populations were mostly manually annotated and typically only compared to MNs, not to other neural sub-types within the developing spinal cord. Thus, the relationship of these *in vitro* engineered cells to their *in vivo* targets remains unclear.

Aiding Capybara analysis of these MN differentiation and programming protocols is a recent single-cell transcriptome atlas detailing the embryonic development of mouse spinal cord populations, surveying the cervical and thoracic spinal cord, spanning five embryonic time points, from E9.5 to E13.5 (Delile et al., 2019). This 38,976-cell spinal cord atlas comprises roof plate (RP), floor plate (FP), and the mantle zone containing distinct classes of neurons which are derived from 11 transcriptionally distinct ventricular zone progenitor cell domains, occupying defined positions within the neural tube. A dorsal-ventral signaling gradient established between the RP and FP directs this regionalization (Alaynick et al., 2011; Briscoe and Small, 2015; Le Dréau and Martí, 2012); (**Figure 4B)**. Altogether, this high-resolution reference encompasses a total of 118 cell types and states, including non-neural cell types surrounding the developing spinal cord, such as blood, mesoderm, and neural crest (Delile et al., 2019).

Under these circumstances, where the developing spinal cord single-cell atlas serves as an ideal reference for our analysis of MN generation, we bypass the initial tissue-level classification step of the Capybara pipeline. Our core interest here is to assess the final identity of MNs produced by these different protocols, rather than characterizing their differentiation trajectories or generation of off-target states. In this instance, the spinal cord atlas, together with an ESC reference from the MCA (Han et al., 2018), serves our objective. Using this combined reference, Capybara discretely classifies 72.0% of cells (n = 3,397/4,719 cells) across the MN programming protocols. 28.0% of cells harbor multiple identities, and 0% of cells remain unclassified. For both the standard and direct programming protocols, Capybara identifies cells mapping to four major embryonic time points: ESC, E11.5, E12.5, and E13.5 (**Figure 4D, Figure S3A**). As expected, the undifferentiated, starting cell population (Day -1/0) is significantly enriched for ESCs (DP: *P* = 0, SP: *P* =0, randomized test; **Figure 4D, Figure S3A**).

Broadly comparing both protocols, ventricular zone neural progenitors are present across both strategies, peaking at early (Day 4/5) and decreasing in late (Day 11/12) stages. These progenitors mainly correspond to developmental stages E12.5 and E11.5 in the standard and direct protocols, respectively (**Figure S3A**). A range of dorsal-ventral progenitor identities is found in the standard protocol, whereas a more restricted progenitor positional diversity is a feature of the direct protocol (**Figure 4C**). In terms of broad neural identity, neuron production peaks at the early stage of the standard protocol (43.9% of Day 5), then subsequently decreases (10.1% of Day 12). In contrast, neural identity is gradually acquired in the direct protocol, with the largest percentage of neurons present in the late timepoint (63.8% of Day 11; **Figure 4C**). This observation is in agreement with previous findings that the largest proportion of neurons is identified on Day 5 for the standard protocol (23%) and Day 11 for the direct protocol (66%) (Briggs et al., 2017). The neural identities we find here mainly correspond to developmental stages E11.5 and E12.5, in the standard and direct protocols, respectively (**Figure S3A**). Similar to the neural progenitors, a range of dorsal-ventral identities can be distinguished, particularly in the direct protocol.

Focusing specifically on MN identity, we observe peak MN production in the early stage of the standard protocol (13.4%), which later decreases to 3.4%. At this later stage, we also observe increased off-target mesoderm identity (1% on Day 5 to 32.9% on Day 12). These observations are in agreement with previous findings proposing that the MN population in the standard protocol decreased to 9% at the later stage due to dominance by the off-target identities. The direct protocol exhibits different characteristics, where only a small percentage of cells (3% on Day 11) emerge as MNs at the late time point. Although this protocol yields MNs, the vast majority (60.8% on Day 11) represent different classes of spinal cord neurons.

Exploring fate transitions in these protocols, we observe a significant decline in transition scores in the early stages of both strategies (Standard protocol: *P* = 1.6E-9; Direct protocol: *P* < 2.2E-16; Wilcoxon Test; **Figure 4E**). From the early to late stage, transition scores remain stable in the standard protocol (*P* = 0.9686; Wilcoxon Test; **Figure 4E**, Left), whereas in the direct protocol, transition scores significantly decrease (*P* < 2.2E-16; Wilcoxon Test; **Figure 4E**, Right). We posit that these scores reflect the maintenance of progenitor-like states in the standard protocol and the emergence of differentiated neurons later in the direct protocol. We next examined the cells harboring multiple identities. In contrast to the hematopoietic transitions we observed, there are very few transitions representing known developmental progressions, suggesting that the mixed identities we observe in this context arise due to defective cell fate specification (**Figure 4F**). This observation, together with the identification of multiple neuron classes, raises the possibility that dorsal-ventral patterning is deficient or absent in both protocols. In the direct protocol, in particular, we observe a wide range of dorsal and ventral neuron identities. Our observations here are supported by the previous identification of a dorsal neuron population in this protocol (Briggs et al., 2017). However, the full range of identities we describe here was not noted in that study, likely because the expression markers defining these neuron classes largely overlap (Delile et al., 2019).

To further interrogate the positional diversity arising in the direct protocol, we leveraged the spinal cord development atlas to identify spinal cord neuron class signatures, based on the combinatorial expression of defined marker genes (Delile et al., 2019). For each cell of the direct protocol dataset, we calculated enrichment scores for these neuron class signatures, projecting these scores onto the UMAP-plot shown in **Figure 4G** and **Figure S3B**. This analysis shows a significant increase in the enrichment of dorsal and ventral neural cell types as the protocol progresses (p < 2.2E-16; **Figure 4H)**. Additionally, the ventral neurons span broader cell types, beyond motor neurons, in which we distinctly identify enrichment of V2a neurons by Day 11 (**Figure 4G)**. Altogether, the data we present here suggests a failure of dorsal-ventral patterning in MN generation from an ESC state, particularly in the direct protocol, raising the possibility that an additional ventralizing signal could boost MN production. Indeed, considering that regionalization of the spinal cord integrates complex spatial and temporal patterning events (Delile et al., 2019), it is not surprising that current protocols do not precisely recapitulate the required spatio-temporal regulation. Indeed, extrinsic signals from paraxial and lateral plate mesoderm are known to enhance the specification of MNs in development (Jessell, 2000). Thus, it raises the possibility that the off-target mesoderm specification in the standard protocol may promote the acquisition of on-target MN identity. Further work is required to explore these hypotheses, but here, Capybara has pointed to an absence of dorsal-ventral patterning in these protocols, creating opportunities to improve on both their efficiency and fidelity.

### Capybara reveals protocol-specific atrial-ventricular patterning in direct cardiac lineage reprogramming

In the above reprogramming system, TFs are overexpressed in ESCs to direct their fate to a specific outcome from a pluripotent state. In a more extreme, but widely used variation of this approach, TF overexpression directly converts cell identity between fully differentiated cell types (Vierbuchen and Wernig, 2011). For example, the overexpression of three TFs: Gata4, Mef2c, and Tbx5 (GMT), can directly reprogram fibroblasts to cardiomyocyte-like cells (Ieda et al., 2010; Qian et al., 2012; Song et al., 2012; Srivastava and Ieda, 2012). Recent studies have characterized this cell fate conversion in-depth (Liu et al., 2017; Stone et al., 2019; Zhou et al., 2019). For example, single-cell transcriptomics has revealed that cells transit through a critical bifurcation to determine their terminal fate in the first two days of conversion (Stone et al., 2019; Zhou et al., 2019). With Capybara, we investigate the cell type composition changes over a time course of reprogramming to cardiac identity.

We selected a recently published single-cell two-week time course of mouse cardiac reprogramming (Stone et al., 2019). In this study, cardiac fibroblasts were harvested from neonatal mice, followed by the introduction of retroviral-delivered TFs, GMT on Day -1. The cells were then treated with TGFβ inhibitor on Day 0 and Wnt inhibitor on Day 1. Day 14 cells were sorted based on the expression of a cardiac reporter gene, α-Myosin Heavy Chain (α-MHC) (Gulick et al., 1991), followed by single-cell profiling to yield a 29,718 cell dataset (Stone et al., 2019). In the first step of Capybara analysis, tissue-level classification suggests the presence of four major tissues in this cardiac reprogramming protocol: neonatal skin, neonatal heart, fetal stomach, and fetal lung (**Figure 5A**, left panel). Additional analysis at the single-cell level using the MCA (Han et al., 2018) narrows this to neonatal skin and neonatal heart (**Figure 5A**, right panel). Two major populations labeled from neonatal skin include macrophages and muscle cells, both of which are mesodermal and resident in the heart (de Soysa et al., 2019).

**Figure 5.**
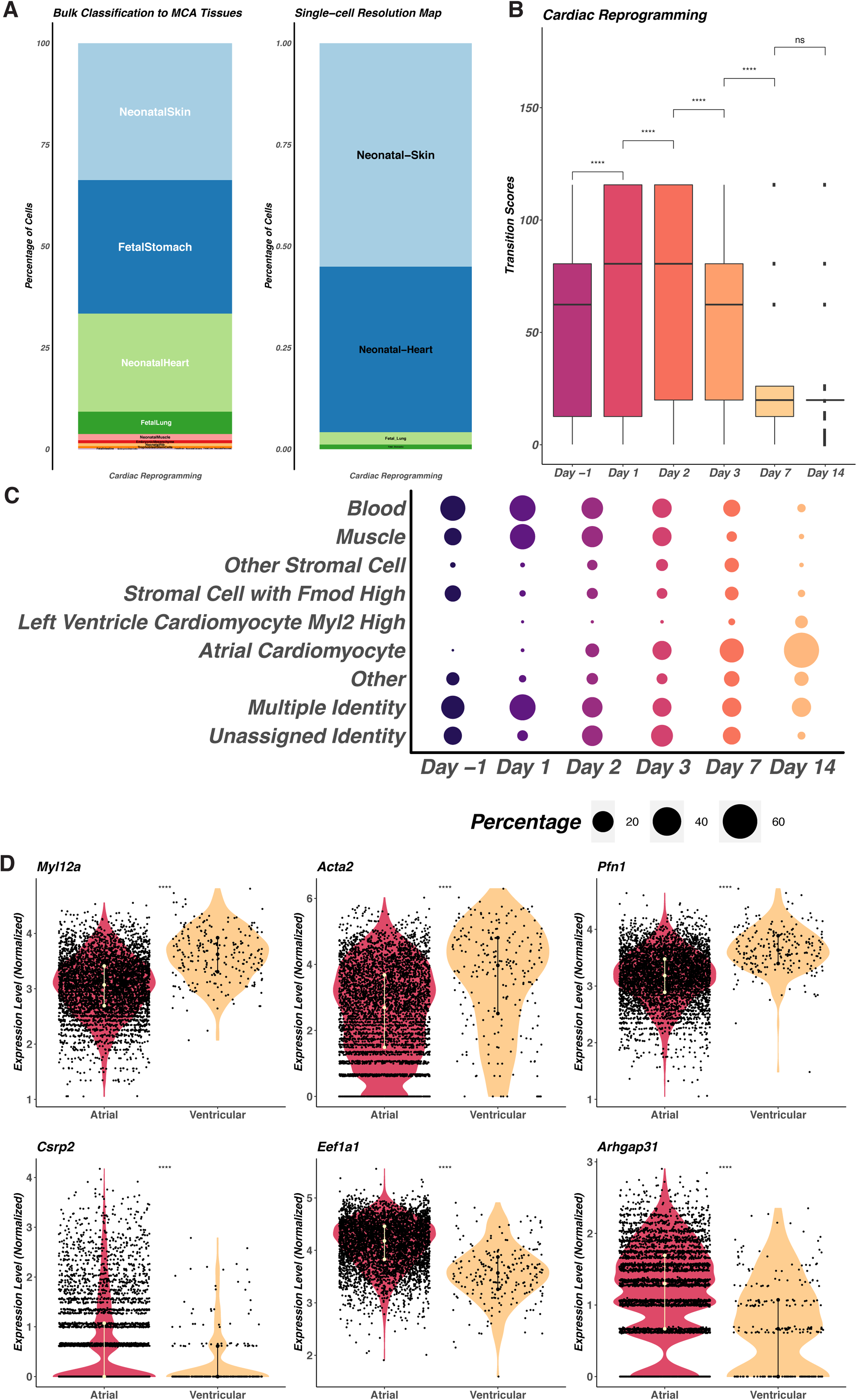
Capybara analysis of cardiac reprogramming. Cardiac fibroblasts were harvested from neonatal mice. Three transcription factors, Gata4, Mef2c, and Tbx5, were retrovirally-delivered into the cells (Day -1), followed by the addition of Wnt inhibitor (Day 0) and TGFβ inhibitors (Day 1). **(A)** Left: In the first step of the Capybara workflow, we perform tissue-level classification of the cardiac reprogramming dataset using ARCHS4, identifying the correlated tissues in the MCA. Major tissues selected in this step include neonatal skin, fetal stomach, neonatal heart, and fetal lung. Right: Tissue composition following higher-resolution classification, refining the major tissues to neonatal skin and neonatal heart. **(B)** Transition scores at each time point (Day -1, Day 1, Day 2, Day 3, Day 7, and Day 14) across the cardiac reprogramming process. A Wilcoxon test was used for significance testing (****: p <= 0.0001, ***: p <= 0.001, **: p <= 0.01, *: p <= 0.05, ns: p > 0.05). **(C)** Cell type composition at each time point. For each timepoint, the size of the dot indicates the percent of the overall population contributed by a single cell type. **(D)** scTransform-normalized gene expression of cardiac markers labeling atrial vs. ventricular cardiomyocytes. A Wilcoxon test was used for significance testing (****: p <= 0.0001, ***: p <= 0.001, **: p <= 0.01, *: p <= 0.05, ns: p > 0.05).

Using the MCA neonatal heart and skin populations as a high-resolution reference, Capybara discretely classifies 65.1% of cells (n = 20,007/30,729 cells) across two weeks of reprogramming to cardiomyocytes. 19.7% of cells harbor multiple identities, and 15.2% of cells are unclassified (**Figure S4A**). Cell type composition analysis reveals a relatively mixed population of starting cells, including blood and muscle (**Figure 5C; Figure S4B**), in agreement with the starting population heterogeneity previously observed (Stone et al., 2019). Cardiac fibroblasts are not explicitly defined in the MCA; however, we identify a stromal cell type (expressing high levels of *Fmod*) that is found only in the neonatal heart that we consider to represent cardiac fibroblasts. We observe a clear signal corresponding to this cell type in the starting population (**Figure 5C**). These non-cardiac populations decrease over time, concomitant with the gradual emergence of cardiomyocytes (**Figure 5C**). In terms of fate transitions, we observe a significant increase in transition scores at Day 1-2 (*P* < 2.22E-16, Day -1 to Day 1; *P* = 2E-7, Day 1 to Day 2; Wilcoxon Test), followed by a progressive decrease at Day 3, which continues to Day 7 (*P* = 2.3E-8, Day 2 to Day 3; *P* < 2.22E-16, Day 3 to Day 7; Wilcoxon Test). These scores imply a period of active fate transitioning in the first two days, followed by a steady fate commitment. This observation echoes previous findings, where the final reprogramming outcome is determined by critical decision-making events in the first 24-48hr following TF expression (**Figure 5B;** (Stone et al., 2019)).

In terms of the properties of the cells generated by this reprogramming strategy, we find that the cardiomyocyte population at Day 14 comprises atrial (75.975%) and left ventricular (7.745%) cardiomyocytes (**Figure 5C; Table S2**). We confirmed the existence of these populations via assessment of region-specific markers (Ventricular markers: *Myl12a, Acta2*, and *Pfn1*; Atrial markers: *Csrp2, Eef1a1*, and *Arhgap31*), demonstrating significant enrichment in the corresponding classified regional cells (*P* < 2.2E-16, Wilcoxon Test**; Figure 5D**). Previous TF-mediated reprogramming studies have reported the generation of mostly atrial-like cardiomyocytes (Efe et al., 2011). From the same group, treatment with small molecules, including a TGFβ inhibitor (SB431542) and Wnt agonist (CHIR99021), during the process reveals a shift from atrial-like to ventricular-like cardiomyocytes (Wang et al., 2014), suggesting a potential chamber specification effect due to changes in Wnt and Bmp signaling pathways, which are known to play a key role in the specification of atrial and ventricular lineages (Bruneau, 2013). In reprogramming with GMT only, mostly ventricular phenotypes were detected, while other subpopulations remain unclear (Ieda et al., 2010). In the reprogramming time course we analyze here, a TGFβ inhibitor (SB431542) and Wnt inhibitor (XAV939) were applied (Stone et al., 2019), but chamber-specific gene expression was not characterized. We hypothesize that the small molecules deployed in this protocol play a role in regionalization, resulting in a population of more atrial cardiomyocytes in this reprogramming paradigm. Here, similar to MN programming, Capybara can capture critical regionalization dynamics, again suggesting potential new strategies to tailor reprogramming outcome.

### Fibroblast to induced endoderm progenitor reprogramming generates a post-injury cell type

Next, with Capybara, we explore another direct reprogramming process that produces a relatively uncharacterized cell identity with no presently known *in vivo* correlate. In this protocol, mouse fibroblasts are reprogrammed to ‘induced endoderm progenitors’ (iEPs), via overexpression of two TFs, Hnf4α, and Foxa1 (**Figure 6A;** (Morris et al., 2014; Sekiya and Suzuki, 2011)). This protocol was originally developed to generate hepatocytes (Sekiya and Suzuki, 2011), yet our subsequent bulk cell type classification and functional analyses revealed that these cells also harbor intestinal potential (Guo et al., 2019; Morris et al., 2014). More recently, using our single-cell resolution lineage tracing approach, CellTagging (Kong et al., 2020), we have identified a bifurcation in this conversion protocol, with one path yielding successfully reprogrammed iEPs, and a path leading to a dead-end state that resembles the original fibroblast population (Biddy et al., 2018; Kamimoto et al., 2020). Based on the hepatic and intestinal potential of iEPs, we have previously suggested that these reprogrammed cells resemble a progenitor-like state (Morris, 2016), although an *in vivo* correlate has remained elusive. To test this hypothesis, we apply Capybara to construct a comprehensive composition map of these cells at different stages of reprogramming, from our previous 85,010 cell lineage tracing time course (Biddy et al., 2018).

**Figure 6.**
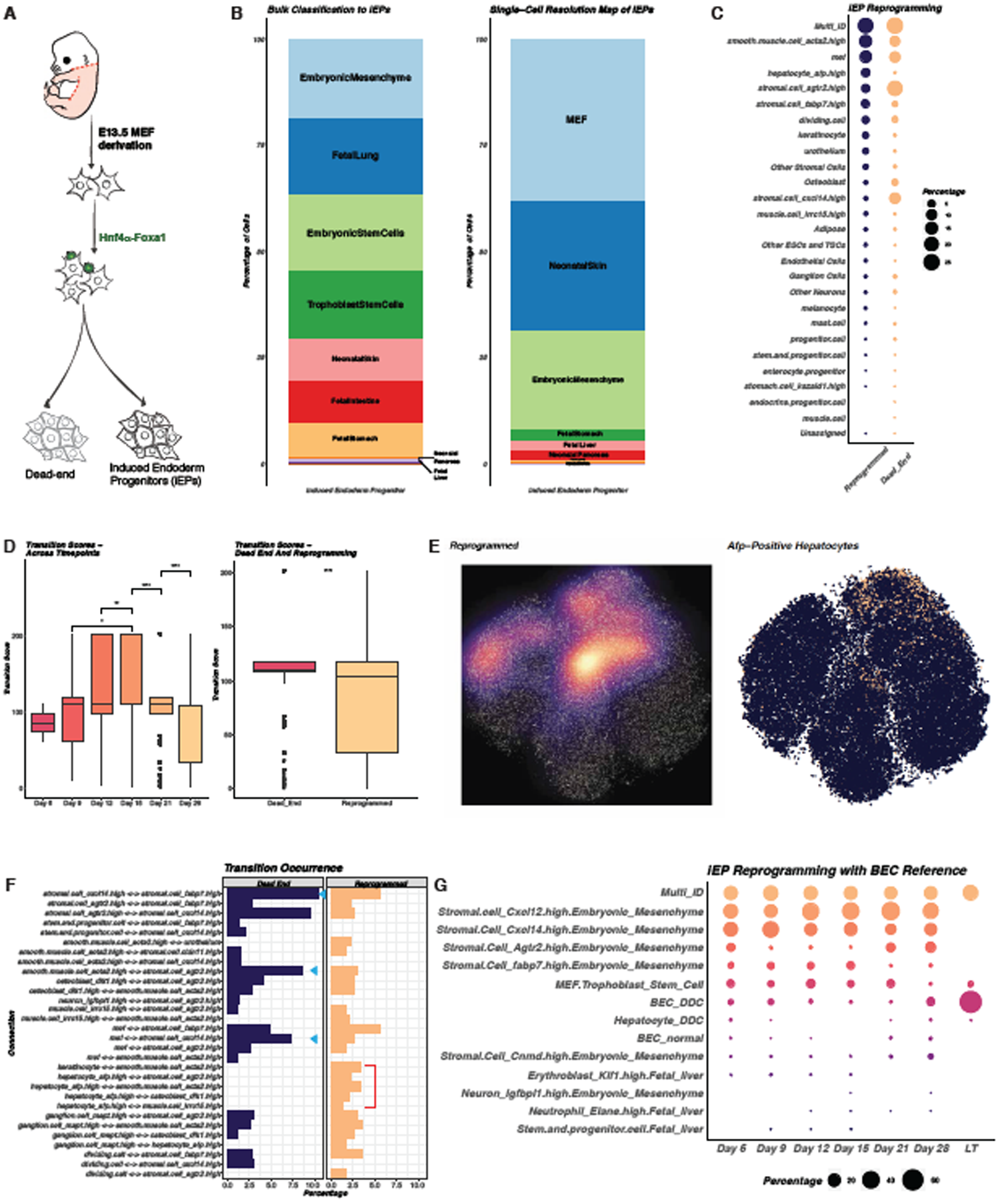
Capybara analysis of fibroblast to induced Endoderm Progenitor (iEP) Reprogramming. **(A)** Schematic of MEF to iEP reprogramming. MEFs are derived from E13.5 mouse embryos. To initiate reprogramming, we overexpress two transcription factors, Hnf4α and Foxa1. Previously, we collected samples from multiple time points during this 28-day reprogramming process and identified two trajectories, one leading to successfully reprogrammed cells, and one path leading to a dead-end (Biddy et al., 2018). **(B)** Left: In the first step of the Capybara workflow, we perform tissue-level classification of the iEP reprogramming dataset using ARCHS4 identifying the correlated tissues in the MCA. Major tissues selected in this step include embryonic mesenchyme, fetal lung, embryonic stem cell, trophoblast stem cell, neonatal skin, fetal intestine, fetal stomach, neonatal pancreas, and fetal liver. Right: Tissue composition after higher-resolution classification, refining the tissues to MEF neonatal skin, embryonic mesenchyme, fetal stomach, fetal liver, and neonatal pancreas. It’s worth noting that in the MCA, MEFs are captured in the trophoblast stem cell tissue type. **(C)** Cell type composition for each reprogramming trajectory. For each timepoint, the size of the dot indicates the percent of the overall population contributed by a single cell type. **(D)** Transition scores plotted for each time point (Left panel; Day 6, Day 9, Day 12, Day 15, Day 21, and Day 28) and transition scores plotted for each trajectory (Right panel; Dead-end vs. Reprogrammed). **(E)** Left: density plot showing cells on the reprogramming trajectory. Right: Cells receiving the ‘*Afp-*expressing hepatocyte’ classification, accounting for 6.8% of cells belonging to the reprogrammed trajectory. **(F)** Comparison of frequencies observed of each transition on the dead-end and reprogrammed trajectories. Blue arrows mark the transitions enriched in the dead-end. The red bracket marks the transitions that are only present in cells on the reprogramming trajectory. **(G)** Cell-type composition of each time point, including the 28-day reprogramming time course, as well as long-term cultured iEPs. These classifications are based on our customized reference containing normal or post-injury BECs and hepatocytes. For each time point, the size of the dot indicates the percent of the overall population contributed by a single cell type.

From initial tissue-level classification, we identify the potential involvement of 9 tissues (with at least 0.5% of cells associated with each tissue), in which embryonic mesenchyme covers the largest proportion of cells. Other tissues include several endodermal types, such as fetal lung, intestine, stomach, pancreas, and liver, as expected for iEPs. ESCs and trophoblast stem cells (TSC), also appeared in this initial correlation map (**Figure 6B**, left panel). We establish a high-resolution reference panel containing a total of 73 cell types across 9 tissues using the MCA (Han et al., 2018). Analysis at this higher resolution narrows these tissues down to 6 major types, including mouse embryonic fibroblasts MEFs (contained within the TSC group in the MCA), neonatal skin, embryonic mesenchyme, fetal stomach, fetal liver and neonatal pancreas (Figure 6A, right panel). Using this reference, Capybara classified 76.6% of the 85,010 reprogramming cells into 47 cell types, with 23.4% of cells sharing multiple identities and 0.032% cells unassigned. The most abundant cell type corresponds to MEFs, representing 30.2% of the cells in total, of which 50% are from Day 0 and Day 3 gradually decreasing to less than 2% at Day 21-28. The other major cell types identified are mesenchymal species, such as stromal cells and smooth muscle cells, in addition to epithelial cell types, such as hepatocytes, and enterocyte progenitors (**Figure S5B**). Across the time course, we observe gradual expansion of epithelial-like cells from 0% at Day 0 (n = 0/11,816 cells) to 10.3% at Day 28 (n = 2,115/20,532 cells), corresponding to cells that have initiated reprogramming. We next investigate the cells with multiple identities, marking transitions during the process. Here on, we focus specifically on a subset of 48,515 cells from Day 6 to Day 28, where CellTag lineage information is enriched, allowing the definition of reprogrammed and dead-end trajectories.

Analyzing cell fate transitions, we observe the highest transition scores between Days 6-15 (*P* = 0.015, Wilcoxon Test; **Figure 6D**, Left), decreasing at Days 21-28. Cells on the reprogramming trajectory receive significantly lower transition scores than the dead-end cells that fail to convert fully (*P* < 2.2E-16, Wilcoxon Test; **Figure 6D**, Right). Interestingly, the variance in the dead-end is smaller than in the reprogrammed population, indicating higher heterogeneity in reprogrammed cells. In support of this, examination of the major compositions of each trajectory reveals eight cell types enriched along the reprogrammed path while two cell types are enriched in the dead-end (**Figure 6C**). In terms of potential transitions, in the earlier phases of conversion, we observe transitions, particularly between MEFs and stromal cells (29.3% of Day 3 transitions; **Figure S5B**). Between Days 15 and 21 of the time course, we see a dramatic shift in the landscape of these transitions, with transitions between stromal cells and smooth muscle, and several cell types to *Afp* expressing hepatocytes (**Figure S5B**). This transition shift coincides with a clear bifurcation at Day 21 toward reprogrammed and dead-end states (Biddy et al., 2018). Here, we only observe transitions to a hepatic state on the reprogramming trajectory, whereas the dead-end path is dominated by stromal transitions, in line with the known regression of these cells back toward a fibroblast-like state (**Figure 6F**); (Biddy et al., 2018; Kamimoto et al., 2020).

Finally, from these analyses, we noted that relatively few reprogrammed cells receive an *Afp*-positive hepatocyte identity (6.8% of reprogrammed cells; **Figure 6E**), whereas a relatively high percentage of cells harbor multiple identities (24.3% of reprogrammed cells). This observation raises the possibility that an *in vivo* correlate of iEPs may not entirely resemble cells contained within the MCA, which largely represents cells in homeostasis. Here, we speculate that multiple identity cells may represent progenitor-like cells that are not present within the MCA, such as facultative stem/progenitor cells that emerge in response to tissue damage. Indeed, our parallel study suggests a role for the Hippo signaling effector Yap1 in the generation iEPs (Kamimoto et al., 2020), in a process resembling the response of cells in the injured liver, where Yap1 is known to play a role in regeneration (Pepe-Mooney et al., 2019), as well as colonic epithelium (Ayyaz et al., 2019; Yui et al., 2018). Considering these observations, in addition to the known potential of iEPs to functionally repair the liver and colon (Guo et al., 2019; Morris et al., 2014; Sekiya and Suzuki, 2011), we hypothesized that iEPs might represent a regenerative epithelial cell type.

To test this hypothesis, we built a high-resolution reference to include MEFs, fetal liver, and embryonic mesenchyme from the MCA. We also integrated recent single-cell profiling from normal and regenerative liver epithelium, which includes two main regenerative cell types: hepatocytes and biliary epithelial cells (BECs) (Pepe-Mooney et al., 2019). Using this reference, we apply Capybara to the time course as well as long-term cultured iEPs that are the source of expanded cells for our previous intestinal transplant studies (Guo et al., 2019; Morris et al., 2014). Classification of the time course iEPs reveals a large proportion of stromal cells across reprogramming, consistent with our above classification using a systematically constructed reference. This demonstrates that the integration of normal and regenerative liver epithelium does not bias the classification. In the population of iEPs at Day 28 in culture, Capybara classifies 7.5% of cells as post-injury BECs, 1.2% as normal BECs, and 0.6% as post-injury hepatocytes. Additionally, Capybara reveals that 67% of our expanded iEPs are discretely classified as post-injury BECs. 29.5% of cells harbor multiple identities, most of which connect a fibroblast state to the post-injury BEC (**Figure 6G**). Evaluation of injured BEC marker expression in the expanded iEPs shows significant enrichment in cells classified as BECs after injury (*P* = 0.00047 *Cyr61*, 9.1E-6 *Klf6*, 2.2E-10 *Krt18*, 0.00065 *Krt8*; **Figure S6A**). Further differential gene expression analysis on the classified cells reveals a set of differentially expressed genes, including *Tuba1b, Stmn1, Rrm2, Birc5*, and *Hmgb2*, that were previously reported to mark proliferative intestinal stem cells post-irradiation (**Figure S6B**) (Ayyaz et al., 2019). We speculate that iEPs might exist as a mixed population of intestinal and liver regenerative cell types, in line with their abilities to engraft damaged intestine and liver (Guo et al., 2019; Morris et al., 2014; Sekiya and Suzuki, 2011). Together, these results highlight the application of Capybara to a relatively uncharacterized reprogramming protocol, incorporating this analysis with our previous lineage tracing studies to offer mechanistic insights into reprogramming, and to identify potential *in vivo* correlates of iEPs, warranting further study.

## Discussion

Here, we have developed and validated Capybara, an unsupervised method to quantitatively assess cell identity and fate transitions. In contrast to current approaches to annotate cell identity, we designed Capybara to interrogate the gradual transition of cell identity during cell differentiation, reprogramming, and disease progression. We benchmark the performance of Capybara against other existing classifiers and demonstrate its efficacy to annotate cells and identify critical transitions within a well-characterized differentiation hierarchy, hematopoiesis. Our application of Capybara to a range of reprogramming strategies reveals previously uncharacterized regional patterning and identifies a putative *in vivo* correlate for an engineered cell type that has remained undefined. These findings prioritize interventions that will potentially increase the efficiency and fidelity of these cell reprogramming strategies, showcasing the utility of Capybara to dissect cell identity across a range of biological systems.

A unique feature of Capybara is the measurement of cell identity as a continuum, and it’s statistical framework to identify cells harboring multiple identities. We build on this concept to define a transition metric, which for each discrete cell identity within a population quantifies the strength and frequency of connections to mixed identity states. A high transition score represents a high information state where identities converge, pinpointing a putative cell fate transition state. This metric is distinct from current approaches that aim to measure cell potential. For example, several platforms use the concept of transcriptional entropy to identify cells with high differentiation potential (Grün et al., 2016; Guo et al., 2016; Teschendorff and Enver, 2017). More recent work has shown that the number of genes expressed per cell can serve as a metric to assess developmental potential (Gulati et al., 2020). A common theme of these approaches is their use of potential measurement to order cells according to their differentiation state.

In contrast, measuring cell potential is not a central aim of Capybara; instead, our metric provides information on whether a cell type is actively transitioning to another identity, revealing those cell states within a system that are actively differentiating, as we demonstrate in our analysis of hematopoiesis. In addition to measuring the overall differentiation ‘flux’ of a biological process, Capybara can also provide information on which identities a cell has the potential to differentiate into, based on shared identities, without requiring their capture within the same experiment. This capability raises several exciting prospects for this analysis. First, we envision that it could help define cell potential in terms of the identities a cell type can give rise to, potentially revealing unanticipated differentiation trajectories. Second, defining transitions may distinguish identity from cell state and determine when cells have reached terminal differentiation. Third, interrogation of transition scores provides information on the overall dynamics of differentiation, helping to pinpoint the timing and nature of pivotal fate decisions. For example, the application of Capybara to reprogramming, in particular, may help identify stochastic and deterministic phases of the process, and help reveal which cell intermediates may be generating off-target identities.

To demonstrate the utility of Capybara to dissect cell identity and fate transitions, we apply it to several fate reprogramming paradigms, an area where the lack of methods to systematically categorize engineered cell identity has represented a challenge. We first analyzed the derivation of motor neurons (MNs) from embryonic stem cells (ESCs) via two distinct routes: growth factor-driven differentiation to mimic embryonic development, and transcription factor (TF) overexpression designed to bypass embryonic states. Indeed, classification and transition scoring shows that TF overexpression produces more terminally differentiated neurons, whereas progenitor states persist in the differentiation approach, in agreement with earlier studies showing that these strategies generally produce developmentally immature phenotypes (Cahan et al., 2014). Focusing on the neural identities produced by both approaches, we observe a range of dorsal and ventral subtypes, suggesting an absence of dorsal-ventral patterning in this system.

Our analysis of cardiac fibroblast to cardiomyocyte shares features with MN programming in terms of regional patterning. We find a bias toward the generation of atrial cardiomyocytes that we posit is due to TGFβ and Wnt signal modulation deployed in this particular protocol (Stone et al., 2019). Based on these findings, and previous work demonstrating that reprogrammed cells more closely resemble target cell types following transplantation into or conversion within an appropriate *in vivo* niche (Guo et al., 2019; Morris et al., 2014; Qian et al., 2012), we posit that strategies combining TF-mediated reprogramming with spatial patterning cues may deliver larger numbers of cells that better recapitulate target cell identity.

Application of Capybara to TF-mediated direct reprogramming of fibroblasts to induced endoderm progenitors (iEPs) provides new observations that may help unlock the identity of this elusive cell type. Previous work has shown that these cells harbor both hepatic and intestinal identity and potential (Guo et al., 2019; Miura and Suzuki, 2017; Morris et al., 2014; Sekiya and Suzuki, 2011). Further characterization of iEPs using CellTagging to assess lineage and identity in parallel revealed a bifurcation in this conversion process, with one path leading to reprogrammed cells and a path to a dead-end fibroblast-like state (Biddy et al., 2018). However, an *in vivo* correlate of these reprogrammed cells has not yet been identified. Here, Capybara classification suggests post-injury biliary epithelial cells (BECs) as a putative *in vivo* correlate of iEPs, warranting further investigation. Together, these observations demonstrate the power of Capybara to enable these highly quantitative observations, suggesting new reprogramming strategies and revealing the nature of target cell identity and potential.

Finally, it is worth noting in our iEP reprogramming analysis, that the absence of relevant target cell types in our current references limited our initial classification. This circumstance highlights the situation where multiple identity cells may represent a stem/progenitor cell type that is absent from available single-cell datasets. Presently, these circumstances require further investigation and the selection/construction of custom atlases. As more diverse single-cell datasets become publicly available, we anticipate that this will support a much broader classification of cell identities. A strength of Capybara to note here is that references can be constructed from a minimum of 30 cells, increasing the likelihood of rare cell type and state capture within our references. A second important caveat: Capybara has a strong dependence on the reference data, not only in terms of cell-type diversity but also in terms of data quality. For example, though gene selection is not required, only genes that intersect between the reference and the sample are used. Thus, for example, the MCA contains expression data for over 8000 genes (Han et al., 2018), while some of our samples contain 15,000 genes. This mismatch effectively leads to downsampling of the test dataset, leading to a loss of some gene expression that could be key to cell-type identification. Again, as technologies improve and cell atlases become more diverse, the accuracy of cell-type classification will increase. In this respect, efforts to standardize, apply quality control, and add additional data modalities to current atlas construction efforts will prove beneficial (Regev et al., 2017). In the future, we anticipate that the addition of multi-omic datasets, such as single-cell ATAC-seq will increase the resolution of Capybara to measure cell identity and fate transitions across a diverse range of biological systems.

## Code availability

Capybara code and documentation are available on GitHub (https://github.com/morris-lab/Capybara).

## Data availability

All source data, including sequencing reads and single-cell expression matrices, are available from the Gene Expression Omnibus (GEO) under accession codes:

- Hematopoiesis Development: GSE72859 (Paul et al., 2015)
- Spinal Motor Neuron Differentiation and Programming: GSE97391 (Briggs et al., 2017)
- Cardiac Reprogramming: GSE131328 (Stone et al., 2019)
- MEFs to iEP Reprogramming Timecourse: GSE99915 (Biddy et al., 2018)
- Long-Term Expanded iEPs: GSE145251
- Human Pancreatic Datasets: GSE84133 (Baron et al., 2016), GSE85241 (Muraro et al. 2016) Single-cell atlas expression matrices are available from:
- Mouse Cell Atlas: <u>Figshare</u> (Han et al.,2018)
- Tabula Muris: <u>Figshare</u> (Tabula Muris Consortium et al., 2018)
- Developing Mouse Spinal Cord Atlas: <u>E-MTAB-7320</u> (Delile et al., 2019)
- Normal and Post Injury Hepatocytes and BECs: GSE125688 (Pepe-Mooney et al., 2019)
- Mouse Gastrulation Atlas: GSE87038 (Pijuan-Sala et al., 2019)
- Bulk transcriptome profiles of all tissues are available from <u>ARCHS4</u> (Lachmann et al., 2018)

## Acknowledgments

We thank members of the Morris laboratory for critical discussions. We thank Barbara Treutlein for sharing the original QP code used in (Treutlein et al., 2016). Thank you also to Colin. This work was funded by National Institute of General Medical Sciences R01 GM126112, and Silicon Valley Community Foundation, Chan Zuckerberg Initiative Grant HCA2-A-1708-02799, both to S.A.M.; S.A.M. is supported by an Allen Distinguished Investigator Award (through the Paul G. Allen Frontiers Group), a Vallee Scholar Award, and a Sloan Research Fellowship.

## Author Contributions

W.K. and S.A.M. conceived the research. W.K. led computational work, assisted by Y.C.F. and supervised by S.A.M. All authors participated in interpretation of data and writing the manuscript.

## Competing Interests

The authors declare no competing interests.

**Correspondence and requests for materials** should be addressed to S.A.M.

## Methods

Capybara code and documentation are available at: https://github.com/morris-lab/Capybara

### Quadratic Programming

Previous studies have measured continuous changes in cell identity using Quadratic programming (QP) (Biddy et al., 2018; Treutlein et al., 2016), where The R package QuadProg was used for the calculation of QP scores. In brief, the underlying assumption is that each single-cell transcriptome profile can exist as a combination of fractional identities from all possible cell types, described as a linear combination of gene expression profiles from different cell types (**Method Figure 1A**). This assumption allows us to model cell identity as a multivariate linear regression problem. For ease of biological interpretation, constraints are placed on the coefficients: they are bound between 0 and 1, and the sum of all coefficients does not exceed 1. These constraints limit the use of least squares estimators in this scenario, while QP is an optimization approach that minimizes a quadratic function under the given linear inequalities or equalities.

Let 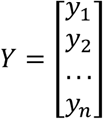 denote the transcriptomic profile of genes *g*_*1*_, *g*_*2*_, …, *g*_*n*_ for a query cell, and *X*_g,t_ denotes the reference dataset of the same set of genes by cell types *t*_*1*_, *t*_*2*_, …, *t*_*m*_. The goal is then to calculate the identity score vector *f*_*t*_, such that the random error ϵ is minimized as described below.

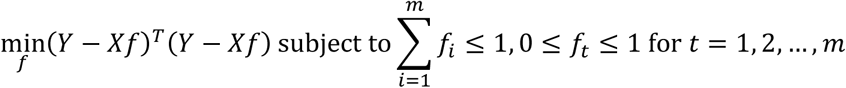

In addition to the fractional identity score matrix and the error term, each cell receives a Lagrangian multiplier, gauging how much the solution is pushed toward the constraints. Applying QP offers a quantitative evaluation of cell identity for each cell.

### Data Processing for QP

Before QP, using raw count matrices, we first perform log-normalization on both the reference and sample dataset. Let *M*_*g,c*_ be the matrix with each row representing a gene, and each column denoting a cell or a cell type. Let *m* denote the number of columns and *n* denote the number of rows. Then, for each column of the matrix, *M*_*\*,c*_,

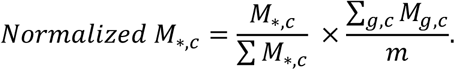

The normalized matrix is then log-transformed with a base of 2, and pseudo-count of 1. The reference dataset undergoes further scaling to ensure that gene expression levels between datasets are comparable. We calculate the scaling factor as the ratio between 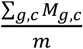 of the reference and sample. Further, we filter the gene list of both matrices to include only those genes shared between the reference and sample.

### Bulk Reference Construction

ARCHS4, a platform that contains most published RNA-seq and ChIP-seq datasets (Lachmann et al., 2018), was mined for bulk RNA-seq data. ARCHS4 obtains raw datasets from the Gene Expression Omnibus, then realigned and processed through a uniform pipeline. Using this data bank, we first filtered the available datasets to retain only poly-A and total RNA-seq data from C57BL/6 mice. We then calculated Pearson’s correlations on every sample pair from the same tissue. The top 90 samples with the highest Pearson’s correlation scores for each of 30 tissues comprised the final bulk reference. For tissues with less than 90 samples, we took the entire sample set and randomly sampled with replacement to include 90 total samples. For the selected 90 samples for each tissue, we calculated the average reads per kilobase per million (RPKM) to build the final tissue-level transcriptome profile, containing a total of 30 tissues. We evaluated the quality of this bulk reference by calculating the identity scores of cells from manually annotated single-cell atlases (MCA; (Han et al., 2018) and Tabula Muris; (Tabula Muris Consortium et al., 2018)) based on this reconstructed reference. We randomly selected 90 cells from each tissue of MCA or Tabula Muris and performed QP using the bulk reference, where we observe high scores when mapping the same tissue between single-cell and bulk datasets (**Method Figure 2**).

### Tissue-Level Classification

A potential concern of using QP to classify single-cells directly is the correlation between similar cell types from different tissues. In this scenario, it could be challenging to tease classification results apart if high similarity to the correct cell type drives the high identity score. Thus, we first perform classification at the tissue level to restrict the number of reference cell types in the downstream analysis, reducing excessive noise and dependencies caused by correlation across tissues in the final single-cell reference. In general, the three primary inputs of this step include the single-cell reference (e.g., MCA), the sample single-cell dataset, and the constructed bulk reference. Using the tissue reference, we calculate QP scores for the single-cell reference as well as the sample, where we obtain two identity matrices. We then compute the Pearson’s correlations of QP scores between each cell from the single-cell reference and each cell from the sample. We use a threshold at the 90^th^ percentile of the correlation matrix to binarize the correlation matrix, where a cell-cell pair with a correlation that is greater than the threshold is marked as 1; otherwise, 0. With the binarized matrix, we count the number of cells in each tissue of the reference mapping to the sample. If there is a significant percentage of reference cells of a tissue (over 70%) mapped, we record the tissue label. We then calculate the frequency of each tissue label in the sample. Tissues with a frequency at least 0.5% sample cells were selected to proceed for further analysis at single-cell resolution. Here, it is worth noting that this tissue-level classification removes most irrelevant tissues but still provides a broad range of tissue types, at which point further downstream analysis removes non-relevant cell types (see ‘Cardiomyocyte Reprogramming Analysis,’ below). Additionally, having prior information regarding the collection tissue can be beneficial to narrow down the tissue selection step, as exemplified by our analysis of hematopoiesis and spinal cord, below.

### Systematic Reference Construction at Single-Cell Resolution

For any tissues selected in the above steps, we leverage available atlases to derive their single - cell transcriptome profiles, to create a high-resolution reference. This reference is assembled based on manual annotation of the specific cell types in the tissue involved. A unique feature of scRNA-seq is dropout - the failure to capture and detect known expressed genes, and other technical variation (Lun et al., 2016). To alleviate the effect of these technical variations, we construct pseudo-bulk references for each cell type of each tissue. We sample 90 cells from each cell type for each tissue. For cell types with more than 90 cells, we calculate Pearson’s correlations between each cell pair. Based on the correlation matrices, we select the most correlated 45 cells to ensure the homogeneity and the least correlated 45 cells to ensure the capture of transcriptional diversity. Cell types that have less than 90 cells but more than 30 cells are sampled with replacement to achieve a total of 90 cells. Summation of the counts of the selected 90 cells is used to construct the final high-resolution reference.

### Empirical P-Value Calculation via Randomized Testing

With the constructed single-cell reference, we apply QP to both the sample and reference single-cell datasets to generate continuous measurements of cell identity (**Method Figure 3A**, left panel). Let *M*_*R*_ denote the identity score matrix of the reference data with a total of *m* cell types and 90 · *m* cells, where *f*_*R,i,j*_ denotes the score of reference cell *i* on cell type *j*. Let *M*_*S*_ denote the identity score matrix of the sample data with a total of *m* cell types and *n* cells, where *f*_*S,i,j*_ denotes the score of sample cell *i* on cell type *j*. We then carry out the following steps to calculate the empirical p-values. 1) For each cell type in *M*_*R*_, we randomly sampled 1000 times and constructed a background density of the identity scores, *D*_*R*_ = [*f* _*resample*,1, …,_, *f*_*resample*,1000_] (**Method Figure 3A**, right top panel). 2) For each score in the identity matrices, we calculate the empirical p-value as following:

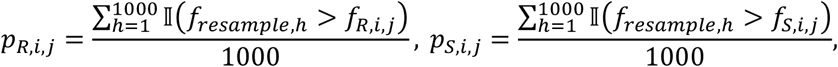

where 𝕀(*) = 1 if (*) is true; otherwise, 𝕀(*) = 0 (**Method Figure 3A**, right bottom panel). 3) Next, we repeat step (1) and (2) for a total of 50 rounds, recording the empirical p-values matrix for each cell of both the reference and the sample. The result of this step includes two lists of p-value matrices: one for the reference and the other for the sample. For each cell, each column of the p-value matrix denotes a cell type, while each row describes each round of 50 (**Method Figure 3B**, left panels).

### Binarization and Classification

From randomized testing, we construct two lists of empirical p-value matrices: one for all sample cells, *P*_*s*_, and the other for all reference cells, *P*_*R*_. Using the list for all reference cells together with its annotation data, we computed a benchmark empirical p-value for each cell type. Specifically, the annotation data contains cell barcodes with their associated annotated cell types (**Method Figure 3B**, right top panel). For each cell *c* and its annotated cell type *t*^0^, we identified the corresponding list of empirical p-values,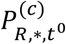. As a result, we construct a possible range of p-values for each cell type, *t*, from which we generate the benchmark values. For each cell type *t*, we eliminate the outlier p-values and select the maximum p-value of the remaining cells as the final benchmark score, *B*_*t*_ = [*B*_*t*1, …_, *B*_*tm*_] (**Method Figure 3B**, middle top panel). Outlier p-values are identified based on the definition of outliers in the boxplot (outside of 1.5x the interquartile range above the third quantile, or below the first quantile).

Next, we move forward to evaluate the sample list. The length of the sample list, *n*, is the number of sample cells. The *n*^*th*^ empirical p-value matrix 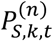 in the list defines empirical p-value for the *n*^*th*^ sample cell belonging to reference cell type *t* under the *k*^*th*^ resampling background, where 1 ≤ *k* ≤ 50. We rank all empirical p-values inside the matrix, from the lowest to the greatest, and break any tie by averaging. The rank-sum for each column *t* of 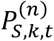 is then calculated, and the cell type with the lowest rank-sum, *t\**, is determined to be the putative identity for cell *c*. We then compare mean 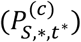 to 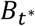, where if 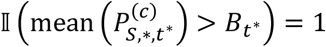, then the identity of this cell is unassigned; otherwise, *t*^*^ is an identity for cell *c*. To identify cells harbouring multiple identities, recapitulating those identities, we perform a pairwise Mann–Whitney U test between the *t\** column and other columns of 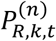. For any cell type *t’* with rank-sum that is not significantly greater than the rank-sum of *t\** (significant level=0.05), we consider *t’* to be one of multiple identities of query *c* along with *t\**. Applying this process to each cell, we generated a binary matrix with 1 = putative identities. Further, we generate a classification table with labeled cell types to each cell barcode (**Method Figure 3B**).

### Transition Scoring

Cells with multiple identities label critical transition states in different trajectories. Building on this concept, we also measure the strength and frequency of connection to the ‘hub’ identities, which provides a metric that we define as a ‘transition score.’ The calculation of transition scores only involves cells with multiple identities. In general, using QP, each cell receives fractional identity scores for different cell types in the reference. Interpreting QP as probabilities of the cell transitioning to each discrete cell identity, we use QP scores as a measure of transition probability. For a cell marked with multiple identities, we consider a transition between the cell to its terminal cell state as events with the transition probability measured by QP scores *P*_*italic*, *j*_, where *i* denotes the cell and *j* denotes the cell state. Therefore, based on information theory, the information of such transition event can be measured as *I*(*transition*) = −log (*P*_*italic*,*j*_). We further consider how much information the terminal cell state has received, which can be defined as:

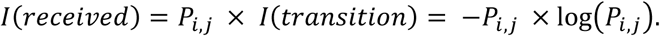

Thus, the total amount of information received for cell state *j* from *n* connected cells can be computed as:

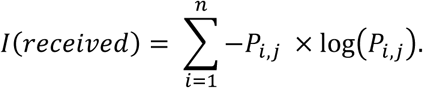

The measurement appears to be similar to Shannon’s entropy, while we note that with each cell independently in transition, probabilities from all events do not necessarily add up to 1, distinguishing it from a measure of entropy. Here, to demonstrate this metric, consider an example as demonstrated in (**Figure 3E**), where Cells 1 to 5 harbor multiple identities connecting Cell State I to III. In this example scenario, the transition score for Cell State II can be calculated as:

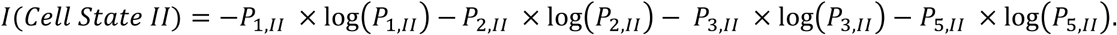

Using such measurement, we incorporate the frequency as well as the likelihood of connection such that high information labels a state as a ‘hub’ state, associated with an abundance of dynamic cell transitions.

### Hematopoiesis Analysis

We obtained the raw hematopoiesis count data from GSE72859 (Paul et al., 2015). The data was processed and clustered using SCANPY (Wolf et al., 2018) and PAGA (Wolf et al., 2019). From processing, we included 3,451 genes in the dataset of 2,730 cells. We first perform tissue-level classification with the bulk reference established using ARCHS4, as described in the previous sections. From this, we identified 3 major relevant tissues: primary mesenchymal stem cells (bone marrow mesenchyme), bone marrow, and bone marrow (c-Kit). Further breakdown of these 3 major tissues using the MCA (Han et al., 2018) resulted in 49 different cell types. We constructed the high-resolution reference using these 49 cell types. From each cell type, 90 cells were selected as described above and saved as the reference single-cell dataset. Followed by preprocessing, we applied QP on the reference and sample single-cell dataset, based on which we further calculate empirical p-values, perform binarization and classification. We projected cells with single identities onto the cluster embedding from PAGA. Cells with multiple identities were isolated, and we extracted the QP scores for their terminal cell identities. We re-assessed these multiple identity cells using their scores. If one of the identities scored near zero (score < 10E-3), we considered such identity as inaccurate and discarded it. In this process, we re-evaluated transitioning cells, retaining only those cells with relatively higher shared identity scores. For a set of mixed identities to be identified as a harbor, we defined a harbor to contain more than 15 cells sharing the same set of multiple identities. We calculated the composition fractions by scaling the QP scores for the terminal identities to a sum of 1. We projected these harbors onto the hematopoiesis lineage tree.

### Gastrulation Transition Score Analysis

We obtained 10x scRNA-seq UMI count data and annotation of mouse gastrulation from GSE87038 (Pijuan-Sala et al., 2019), containing a total of 139,331 cells. The dataset was processed using Seurat (Butler et al., 2018; Satija et al., 2015). We performed classification using all 23 tissues, composed of 361 cell types, in the adult MCA as reference directly (Han et al., 2018). We constructed the high-resolution reference using these annotated cells. Following preprocessing, we generated the continuous identity score measurements for these cells using QP, followed by binarization, and classification. We then performed Capybara transition scoring analysis for each sample, analyzing transition score distributions of each annotated cell type from Pijuan-Sala et al.

### *In Vitro* Spinal Cord Motor Neuron Derivation Analysis

We obtained 10x scRNA-seq UMI count data and annotation of developing mouse spinal cord from https://www.ebi.ac.uk/arrayexpress/experiments/E-MTAB-7320/ (Delile et al., 2019), including a total of 38,976 cells. We removed unannotated cells and built a high-resolution reference for each cell type at each developmental stage (E9.5 to E13.5), resulting in a total of 118 cell types (19 types for E9.5, 26 for E10.5, 26 for E11.5, 25 for E12.5, 22 for E13.5). We also included embryonic stem cells from the MCA (Han et al., 2018). We obtained the InDrop single-cell dataset for *in vitro* spinal cord motor neuron derivation from GSE97391 (Briggs et al., 2017). The *in vitro* datasets were processed and clustered using SCANPY (Wolf et al., 2018) and PAGA (Wolf et al., 2019). From processing, we included 7,860 genes and 7,799 genes in the dataset of 1,984 cells and 2,720 cells in direct programming (DP) and standard protocol (SP), respectively. We analyzed ESCs from each protocol separately. Following preprocessing, we applied QP using the high-resolution reference on 4 datasets, including two ESC populations, DP and SP. Based on the identity score matrices, we calculated the empirical p-value matrices, performed binarization, and classification. Cells with single-identities were separated to calculate the composition in the ventricular zone and mantle zone. The ventricular zone also included the neural crest neurons and mesoderm lineage. Cells with multiple-identities were filtered and refined, as described in the above hematopoiesis section. With the QP scores attached to each identity on each identity in the mixed set, we calculated the transitions scores for the cell states involved, as described in the transition scoring section. We compared transition scores between different timepoint via a one-sided Wilcoxon test.

### Cardiomyocyte Reprogramming Analysis

We obtained The 10x single-cell RNA-sequencing count data from GSE131328 (Stone et al., 2019) containing a total of 30,729 cells. This dataset was processed using Seurat (Butler et al., 2018; Satija et al., 2015) and clustered using UMAP. We used raw data from the filtered cells and genes as input into the Capybara pipeline. We next performed tissue-level classification using ARCHS4 (Lachmann et al., 2018), as described in previous sections, revealing 4 major tissues, including neonatal skin, neonatal heart, fetal stomach, and fetal lung. Further breakdown of these tissues using MCA (Han et al., 2018) contains a total of 57 cell types. We constructed the high-resolution reference using these annotated cells. Following preprocessing, we generated the continuous identity score measures of these cells using QP, based on which we further performed binarization, and classification. We calculated the percentage of each identified cell types in the population. Additionally, we computed the transition scores for the cell states involved in transitions. We performed transition score comparisons using a one-sided Wilcoxon test. We identified region specific markers from MuscleDB (http://muscledb.org/mouse/mRNA/).

### Induced Endoderm Progenitor Analysis

We processed scRNA-seq data of induced endoderm progenitor (iEP) reprogramming, as previously described (Biddy et al., 2018). In brief, Scater was used to normalize (McCarthy et al., 2017) the data across time points and Seurat (Butler et al., 2018; Satija et al., 2015) was used to integrate biological replicates, perform clustering, and visualize cells using *t*-SNE. We performed clone calling using the following workflow: https://github.com/morris-lab/CellTagR. We used previous annotations to define cells on the reprogramming or dead-end trajectories (Biddy et al., 2018). We performed tissue-level classification using ARCHS4 (Lachmann et al., 2018), as described in previous sections, highlighting the involvement of 9 potential tissues, containing a total of 73 cell types. Following construction of the high-resolution reference, we performed preprocessing on the reference and the sample, on which we then applied QP to generate the identity score matrices. Further, we calculated the p-value matrices and performed binarization and classification. We calculated the percent composition of each cell type. Cells with multiple identities were filtered as described in the above hematopoiesis section, except a harbor was defined in this scenario to have more than 30 cells, taking into account the larger population sampled in this dataset. We performed transition scoring, as above.

We obtained scRNA-seq data of biliary epithelial cells (BECs) and hepatocytes, before and after injury, from GSE125688 (Pepe-Mooney et al., 2019). We built a custom high-resolution reference by incorporation of additional tissues from the MCA: fetal liver, MEFs, and embryonic mesenchyme. The long-term iEPs were cultured for 12 months before collection and processing. We had previously used these cells to engraft mouse colon (Guo et al., 2019; Morris et al., 2014). The long-term iEP dataset was processed, filtered, and clustered using Seurat, resulting in a total of 2,008 cells. We then constructed the high-resolution reference panel with 20 cell types and performed preprocessing on the reference and single-cell sample. Application of QP using processed reference and long-term iEP and iEP reprogramming datasets provides us the continuous metric of identity scores, from which we further carried out binarization and classification. We compared gene expression between groups using a Wilcoxon test.

### Long-term iEP culture and single-cell profiling

Mouse Embryonic Fibroblasts were derived from the C57BL/6J strain (The Jackson laboratory: 000664). All animal procedures were based on animal care guidelines approved by the Institutional Animal Care and Use Committee. Mouse embryonic fibroblasts were converted to iHeps/iEPs as in Sekiya and Suzuki (2011). Briefly, fibroblasts were prepared from E13.5 embryos and serially transduced with polyethylene glycol concentrated Hnf4α-t2a-Foxa1, followed by culture on gelatin for 2 weeks in hepato-medium (DMEM:F-12, supplemented with 10% FBS, 1 mg/ml insulin (Sigma), dexamethasone (Sigma-Aldrich), 10 mM nicotinamide (Sigma-Aldrich), 2 mM L-glutamine, 50 mM β-mercaptoethanol (Life Technologies), and penicillin/streptomycin, containing 20 ng/ml hepatocyte growth factor (SigmaAldrich), and 20 ng/ml epidermal growth factor (Sigma-Aldrich), after which the emerging iEPs were cultured on collagen and passaged twice per week for 3 months. For single-cell library preparation on the 10x Genomics platform, we used: the Chromium Single Cell 3′ Library & Gel Bead Kit v2 (PN-120237), Chromium Single Cell 3′ Chip kit v2 (PN-120236) and Chromium i7 Multiplex Kit (PN-120262), according to the manufacturer’s instructions in the Chromium Single Cell 3′ Reagents Kits V2 User Guide. Just prior to cell capture, methanol-fixed cells were placed on ice, then spun at 3000rpm for 5 minutes at 4°C, followed by resuspension and rehydration in PBS, according to (Alles et al., 2017). 17,000 cells were loaded per lane of the chip, aiming to capture 10,000 single-cell transcriptomes. The resulting cDNA libraries were quantified on an Agilent Tapestation and sequenced on an Illumina HiSeq 2500.

## Methods

**Method Figure 1. (A)** The general assumption of quadratic programming (QP). Each single-cell transcriptome is a linear combination of gene expression profiles of different cell types.

**Method Figure 2**. Mapping of single-cell atlases to mined ARCHS4 bulk transcriptome. Left panel: Adult and fetal tissues from Mouse Cell Atlas (MCA; Han et al., 2018). Right panel: Tabula Muris with different technologies (Tabula Muris Consortium et al., 2018)

**Method Figure 3. (A)** Flow chart to calculate empirical p-values. Details can be found in section ‘Empirical P-Value Calculation via Randomized Testing’. **(B)** Flow chart to perform binarization and classification, using example meta data from hematopoiesis. See ‘Binarization and Classification’ for details.

**Method Validation: Benchmarking Capybara**

To assess the efficacy and robustness of Capybara to classify cell identity, here we validate each step and demonstrate its basic functionality. In the first step of the Capybara pipeline, tissue-level classification, accuracy is pivotal as it helps reduce noise from other cell types that are not present in the sample. We evaluate the validity of the tissue reference transcriptome based on the identity scores of annotated single-cell atlases (Han et al., 2018; Tabula Muris Consortium et al., 2018). We randomly selected 90 cells from each tissue of MCA and Tabula Muris using the bulk reference, where we observed higher scores mapping of the same tissue between single - cell and bulk (**Method Figure 2**).

Next, we assess the classification functionality of Capybara. In this step, we use a benchmarking algorithm that was recently developed to compare a range of single-cell classification approaches using an array of publicly available datasets (Abdelaal et al., 2019). Briefly, 10-fold cross validation is performed using various datasets, and predictions from the methods are assessed based on accuracy, median F1 score, proportion of undefined cells, and classification runtime. Based on 5 human pancreatic datasets, the performance of Capybara indicates similar accuracy (rank 5) and median F1 score (rank 5) with reasonable runtime, when benchmarked against 8 other classifiers (**Figure S1B**). In this automatic benchmarking method, 5-fold cross-validation provides a relatively large training set (80%), compared to the test set (20%). A key feature of Capybara is its flexible requirement in terms of training set size. We find that a minimum number of 90 cells sampled from each cell type is required to perform accurate classification. For cell types that have fewer than 90 cells, we require a minimum of 30 cells, from which a 90-cell sample will be drawn with replacement from the pool.

Using this minimum number of cells, we evaluate our performance using the *Tabula Muris* mouse cell atlas (Tabula Muris Consortium et al., 2018). Using Cohen’s Kappa scores and accuracy, we benchmark our method against two other classification approaches, scmap (Kiselev et al., 2018) and SingleCellNet (Tan and Cahan, 2019). As a result, we demonstrate the comparable performance of Capybara with almost perfect agreement (Cohen’s Kappa score > 0.81) in 80% of the tissues (**Figure S2A**). As Cohen’s Kappa score can be sensitive to off-target prediction, we inspect the accuracy of prediction for tissues with low Cohen’s Kappa score, where the percentage of agreement, a major calculation for accuracy, is around 90% (**Figure S2A**).

Moreover, we evaluate the ability of Capybara to support accurate cross-platform classification. Here, we showcase this function using two human pancreatic islet datasets: Baron et al., 2016 (inDrop, reference) and Muraro et al., 2016 (CEL-seq2, test). Cross-platform validation presents the agreement between annotation and classification with a Cohen’s Kappa score of 0.967 considering the cell type annotations shared between the two datasets. Four major off-target cell-type classifications are observed. Investigating these cells, we separate them into two categories: One category includes cell types that are annotated in the test set but not in the reference; the other category includes cell types that are misclassified. In the former, we explore the markers used in the reference for labeling to investigate whether the cells annotated differently in the test set can map to other cell types in the reference. From Capybara, mesenchymal cells in the test are classified as activated stellate cells. Mesenchymal cells take broad definition, potentially including stellate cells in the pancreas. Expression of *PDGFRA* shows significant enrichment in the classified activated stellate cells compared to other groups. Similarly, PP cells are determined to be gamma cells, whose marker genes, including *PPY, SERTM1* and *CARTPT*, show upregulation in the classified cells. We further determine cells with unknown cell types to be ductal cell-like. In the latter category, it is interesting to note that Capybara identifies heterogeneity in the annotated acinar cells. Using three acinar cell marker genes, including *REG3A, PRSS1* and *CPA1*, we cannot isolate expression differences between labeled acinar cells and ductal cells. While, *KRT19*, a ductal cell marker, reveals a significant upregulation in the labeled ductal cells, indicating that *KRT19* might play a key role in classification of ductal cells. Additionally, it is worth notice that the original studies use slightly different marker selection for cell type annotation, potentially leading to mismatch between datasets. Overall, this analysis demonstrates the efficacy Capybara to classify cell identity using cross-platform data sources.

## Supplementary Figure Legends

**Supplementary Figure S1. Evaluation and benchmarking of the Capybara Pipeline. (A)** Correlation heatmap of bulk expression between each pair of the 30 identified tissues from ARCHS4. The bulk expression is the logged reads per kilobase per million (RPKM). The RPKM was calculated as the average of the 90 samples selected for each tissue (See Methods). The color of each square denotes the correlation between tissues, where a lighter color indicates a higher correlation. **(B)** Evaluation of using 5 human pancreatic dataset benchmarking against 8 other classifiers using an established benchmarking pipeline (Abdelaal et al., 2019). This algorithm evaluates four aspects of the algorithm, including accuracy, median F1 scores, mean total time, and unlabeled percentage. Color of each square labels the assessment of each aspect.

**Supplementary Figure S2. Additional benchmarking of the Capybara Pipeline. (A)** Cross-validation using *Tabula Muris* processed with 10x droplet-based or smart-seq2 technologies. In this evaluation, 90 cells were sampled from each cell type of each tissue to construct the high-resolution reference. Sample cells used for reference construction were used as training sets for scmap and SingleCellNet. The remaining cells were used as test samples. Accuracy and Cohen’s Kappa score of the classification on the test sample were calculated against their original annotation. **(B)** Cross-platform validation between Baron et al., 2016 (InDrop) and Muraro et al., 2016 (CEL-Seq2). Left: Proportions of cells mapping between Capybara classification and the original annotation. A lighter color indicates a higher percentage agreement. Right: Log-normalized expression levels of marker genes across classified cell types, calculated using Seurat. The marker genes were identified from Baron et al., 2016. Gamma cells: *PPY, SERTM1, CARTPT*; Activated stellate cells: *PDGFRA*; Acinar cells: *REG3A, PRSS1, CPA1*; Ductal cells: *KRT19*.

**Supplementary Figure S3. Capybara analysis of spinal motor neuron differentiation and programming. (A)** Detailed breakdown of cell types classified in the two protocols to generate motor neurons (MNs) from embryonic stem cells (ESCs). The percentage of each cell type was calculated based on each time point of each protocol. Top: Cell types in the ventricular zone, including mesoderm cell types. Bottom: Cell types in the mantle zone. Left: Standard protocol. Right: Direct programming. **(B)** Projection of enrichment scores of dorsal-ventral neuron identities onto UMAP visualization of the direct programming protocol. Top Left: Projection of time points (Day -1, Day 4, Day 11). Marker genes of each region of the mantle zone were identified from a previously curated gene list (Delile et al., 2019). The enrichment scores were calculated using the *AddModuleScore* function in Seurat, using the curated gene list.

**Supplementary Figure S4. Capybara analysis of cardiac reprogramming. (A)** A breakdown of the cardiac reprogramming into classification subtypes, including single-identity, multiple-identity, and unassigned-identity calls. **(B)** Detailed cell-type breakdown for each time point of cardiac reprogramming (Day -1, Day 1, Day 2, Day 3, Day 7, Day 14). For each timepoint, the size of the dot indicates the percent of the overall population contributed by a single cell type.

**Supplementary Figure S5. Capybara analysis of fibroblast to induced Endoderm Progenitor (iEP) Reprogramming. (A)** Detailed cell-type breakdown for each time point of the entire reprogramming time course (Day 0, Day 3, Day 6, Day 9, Day 12, Day 15, Day 21, Day 28). For each timepoint, the size of the dot indicates the percent of the overall population contributed by a single cell type. **(B)** Frequency of each transition at each time point, starting from Day 3 to Day 28, where a dramatic shift in the landscape of these transitions occurs between Day 15 and Day 21.

**Supplementary Figure S6. Marker analysis of MEF to iEP Reprogramming using a customized reference with normal and post-injury hepatocytes and biliary epithelial cells (BECs). (A)** Log-normalized BEC marker gene expression for classified cell types. The marker genes are identified from Pepe-Mooney et al., 2019. **(B)** Log-normalized proliferative intestinal stem cell marker gene expression for classified cell types. The marker genes are identified from Ayyaz et al., 2019. A Wilcoxon test was used for significance testing.

