## Supplementary Figures for "Capybara: A computational tool to measure cell identity and fate transitions"

### Supplementary Figure S1

**A**

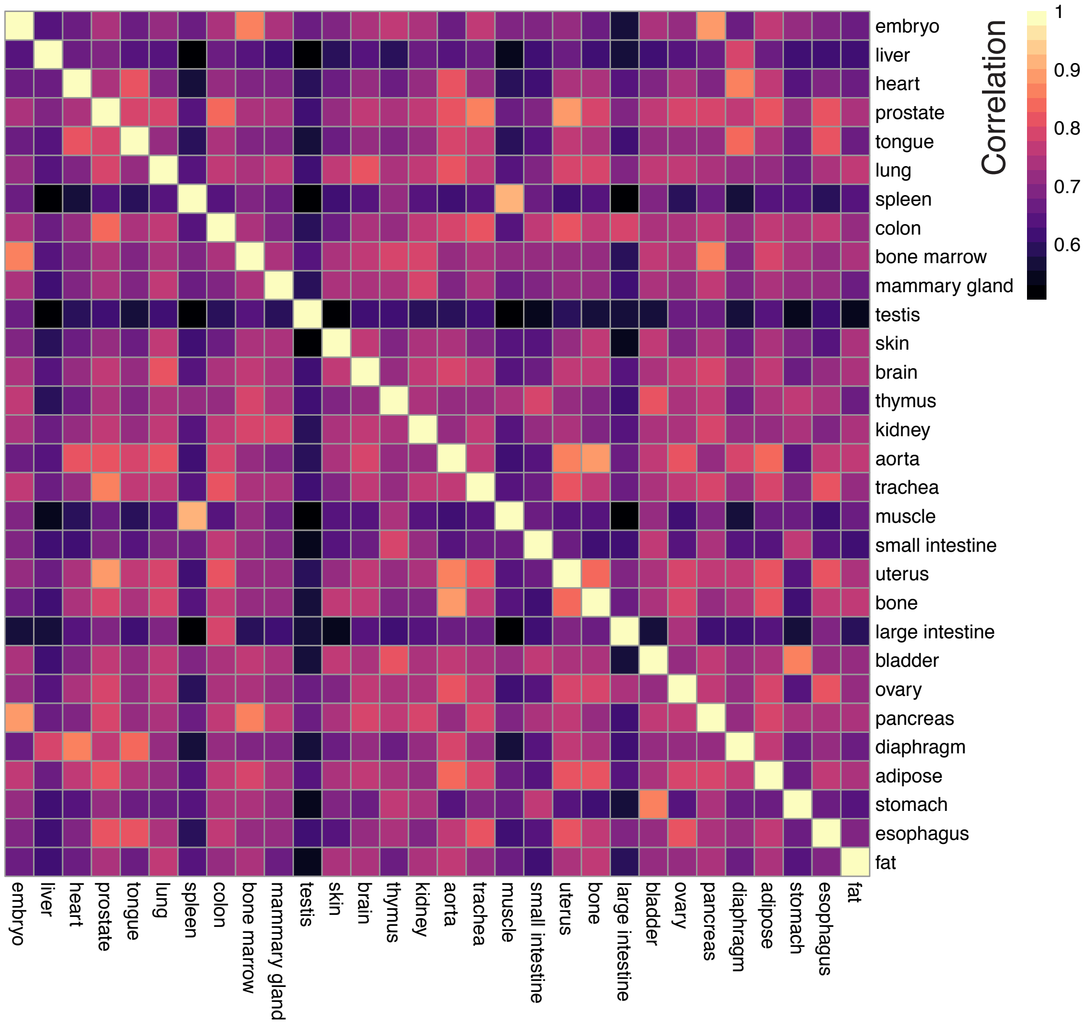


#### B *Pancreatic Dataset Evaluation*

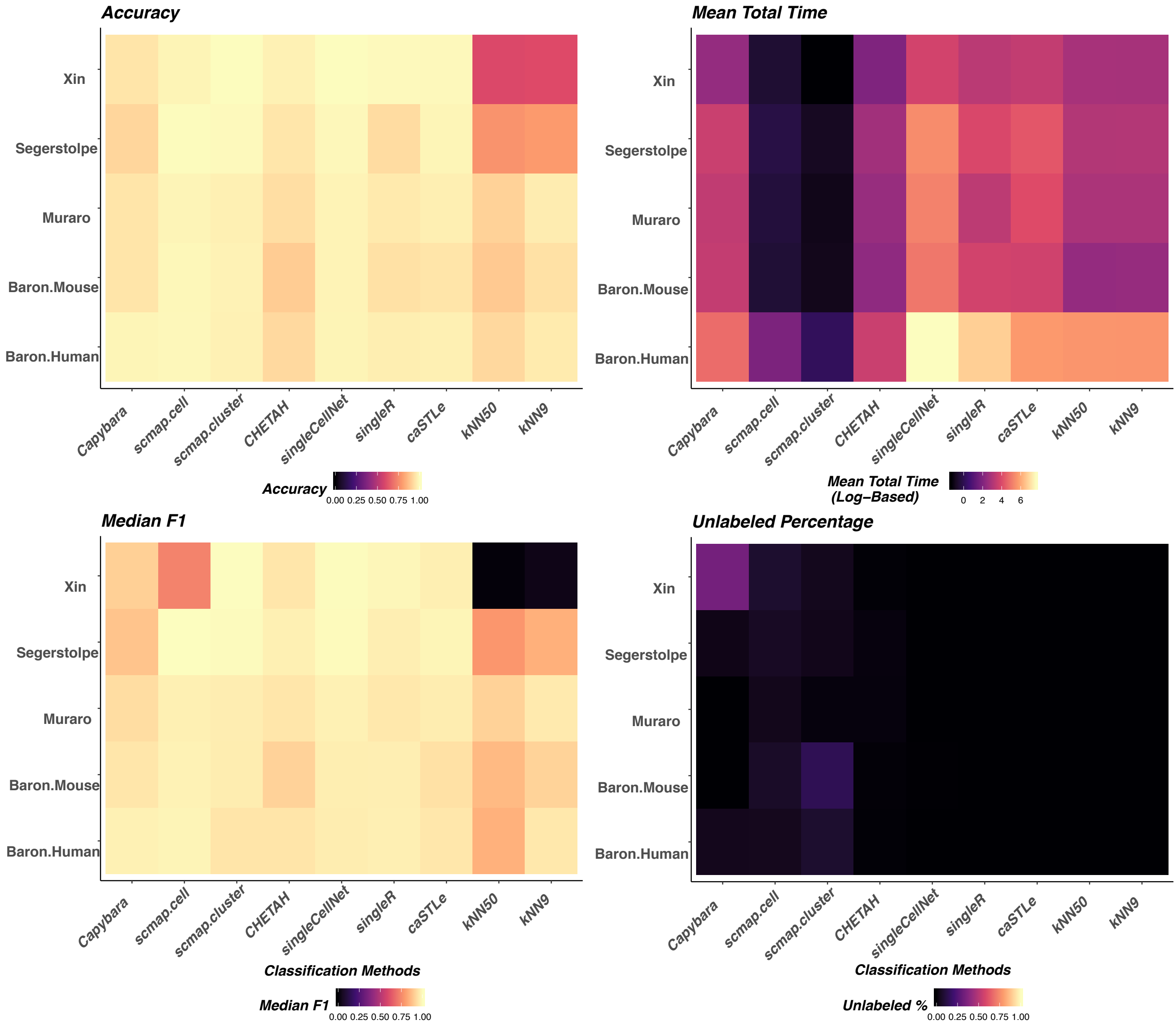

### Supplementary Figure S2

#### A. Cross-Validation Using *Tabula Muris*

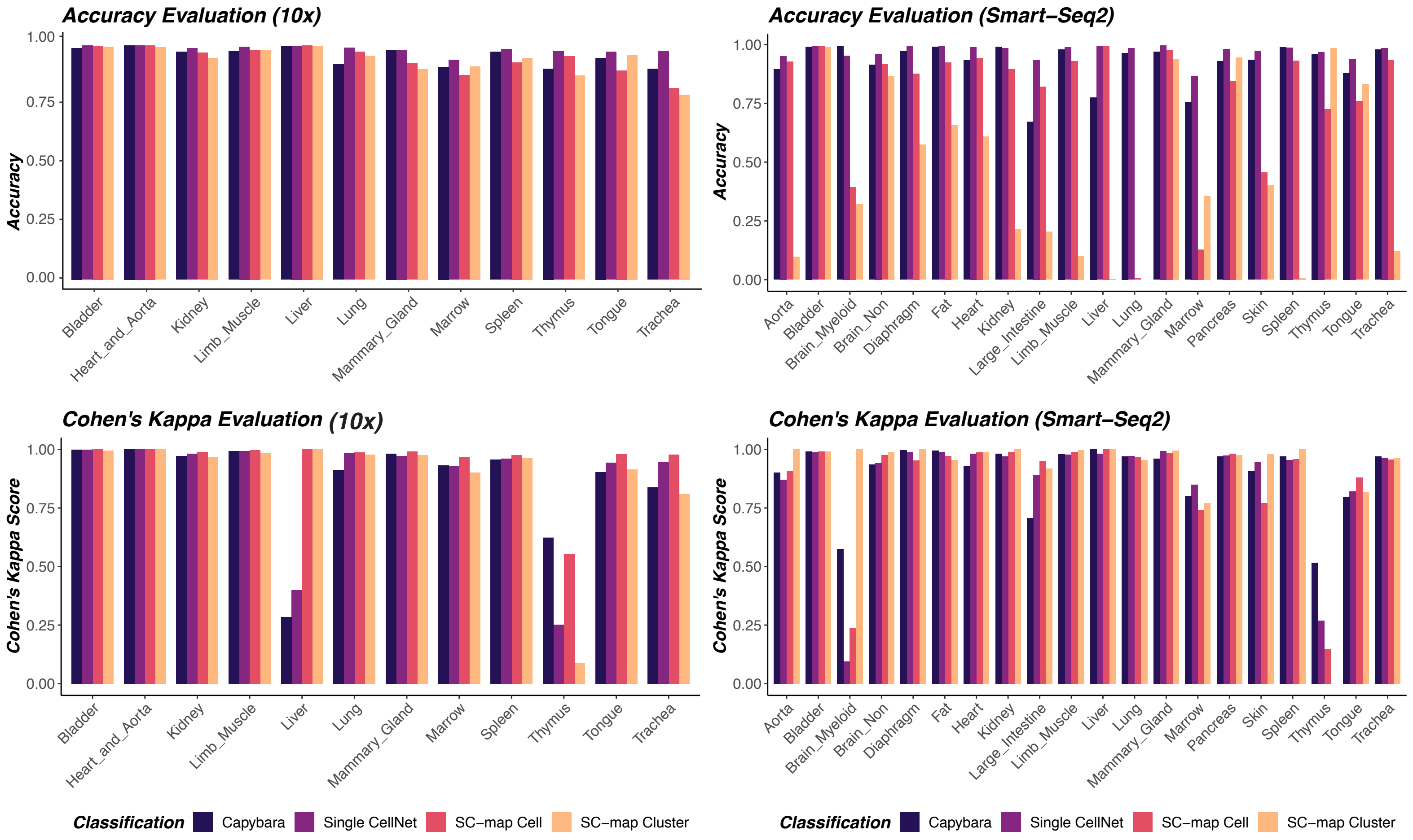


#### B. Cross-Platform Validation between Baron *et al* and Muraro *et al*

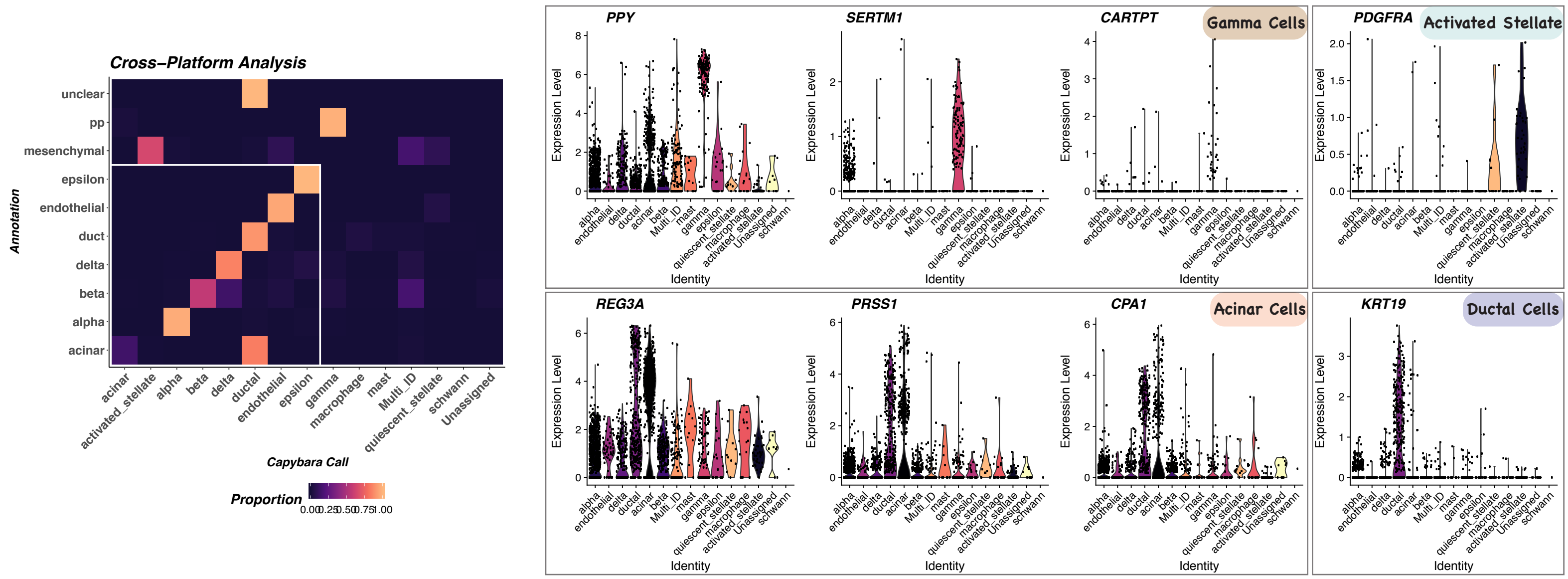

### Supplementary Figure S3

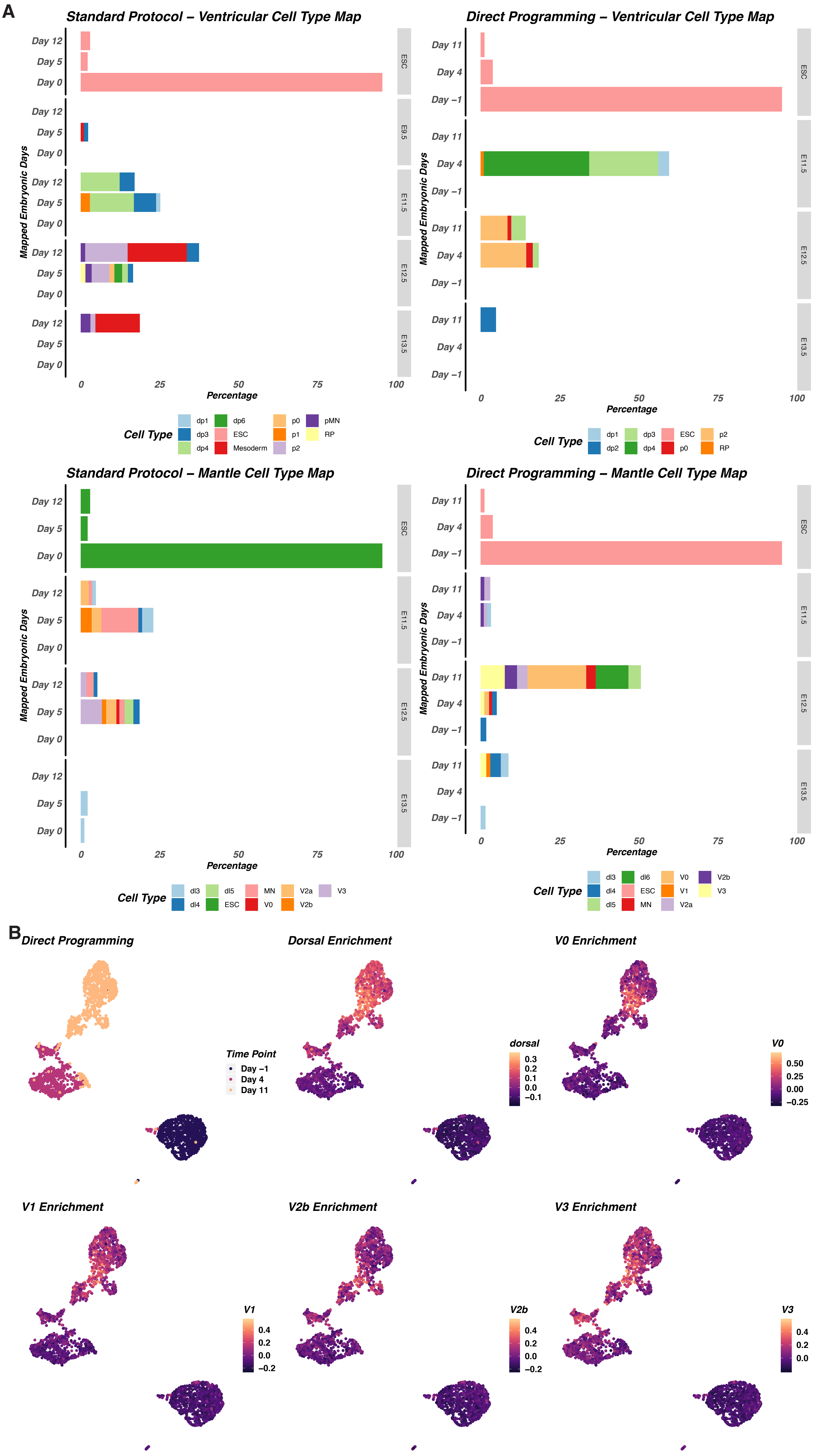

### Supplementary Figure S4

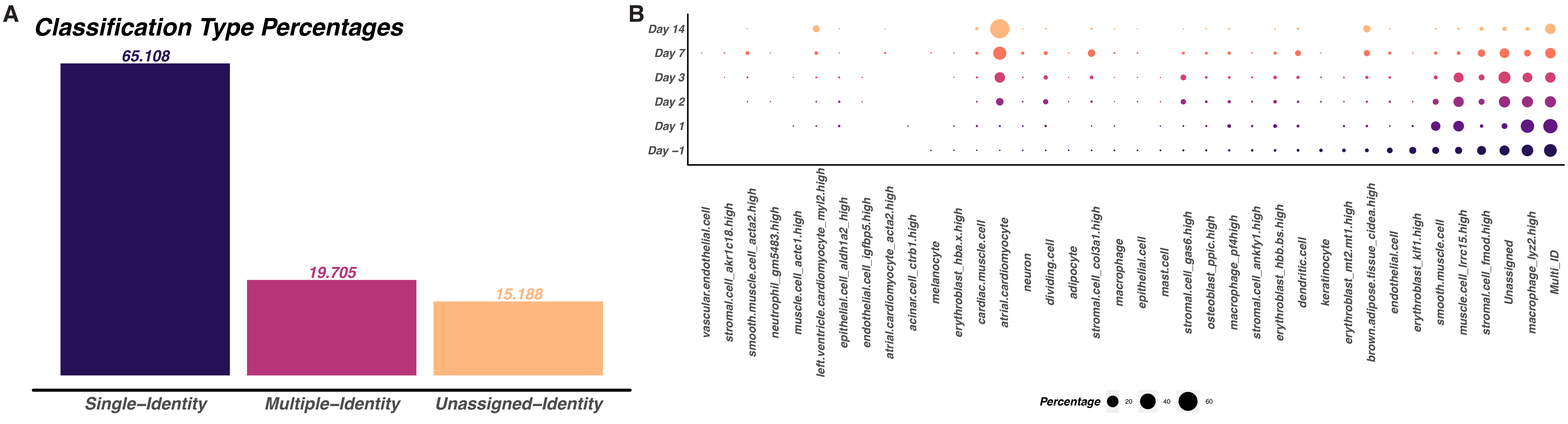

### Supplementary Figure S5

**A**

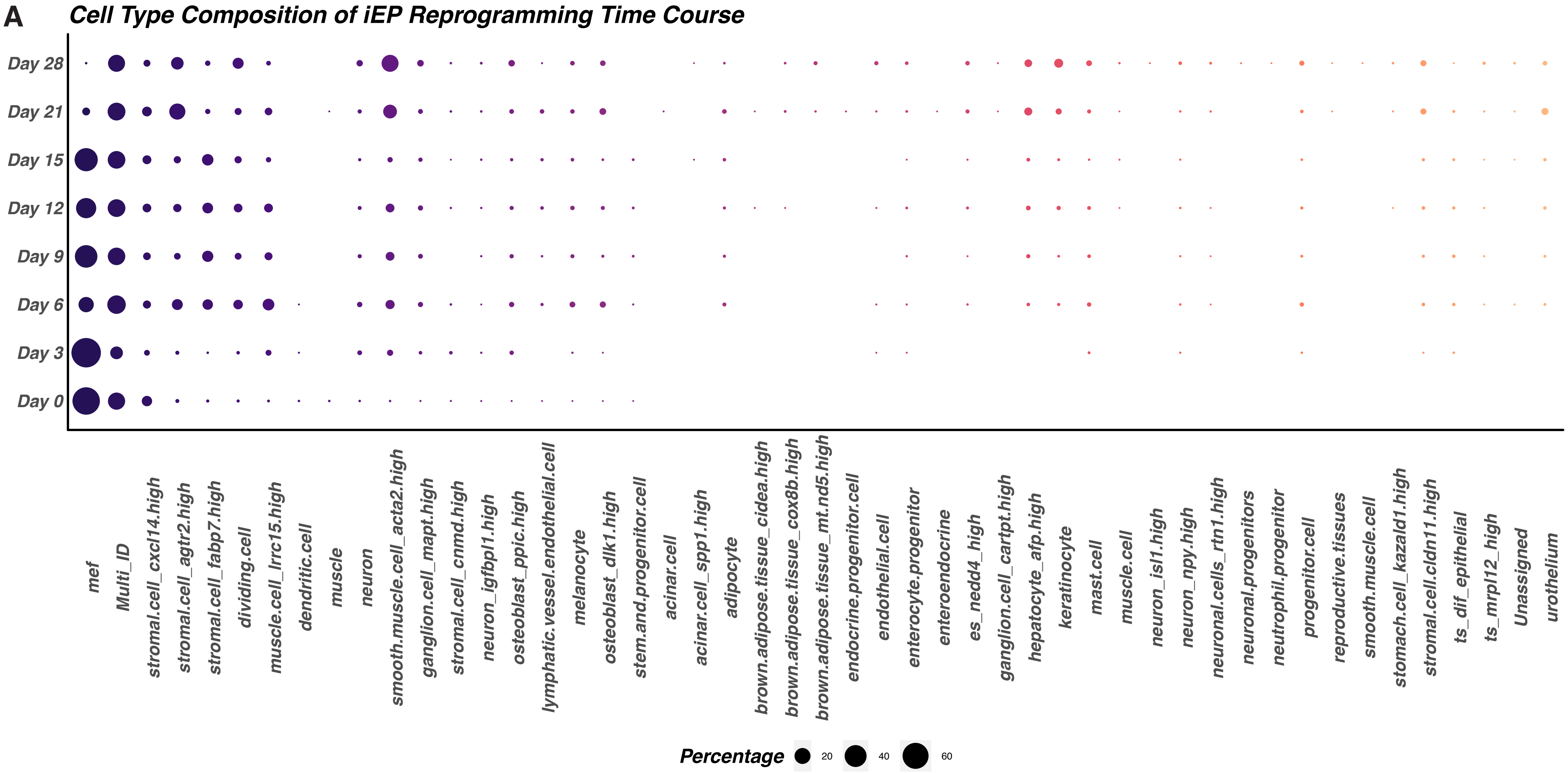


**B**

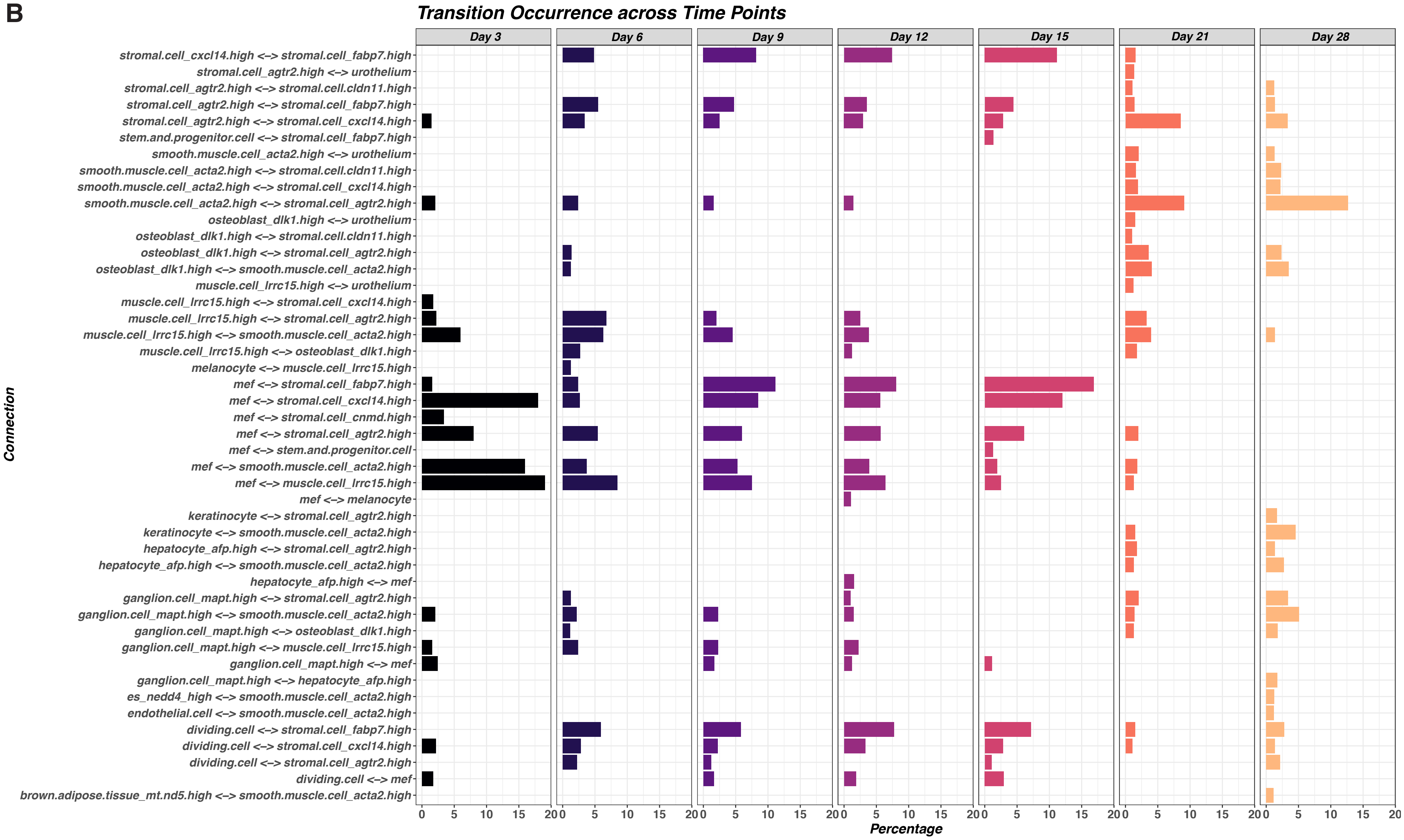

### Supplementary Figure S6

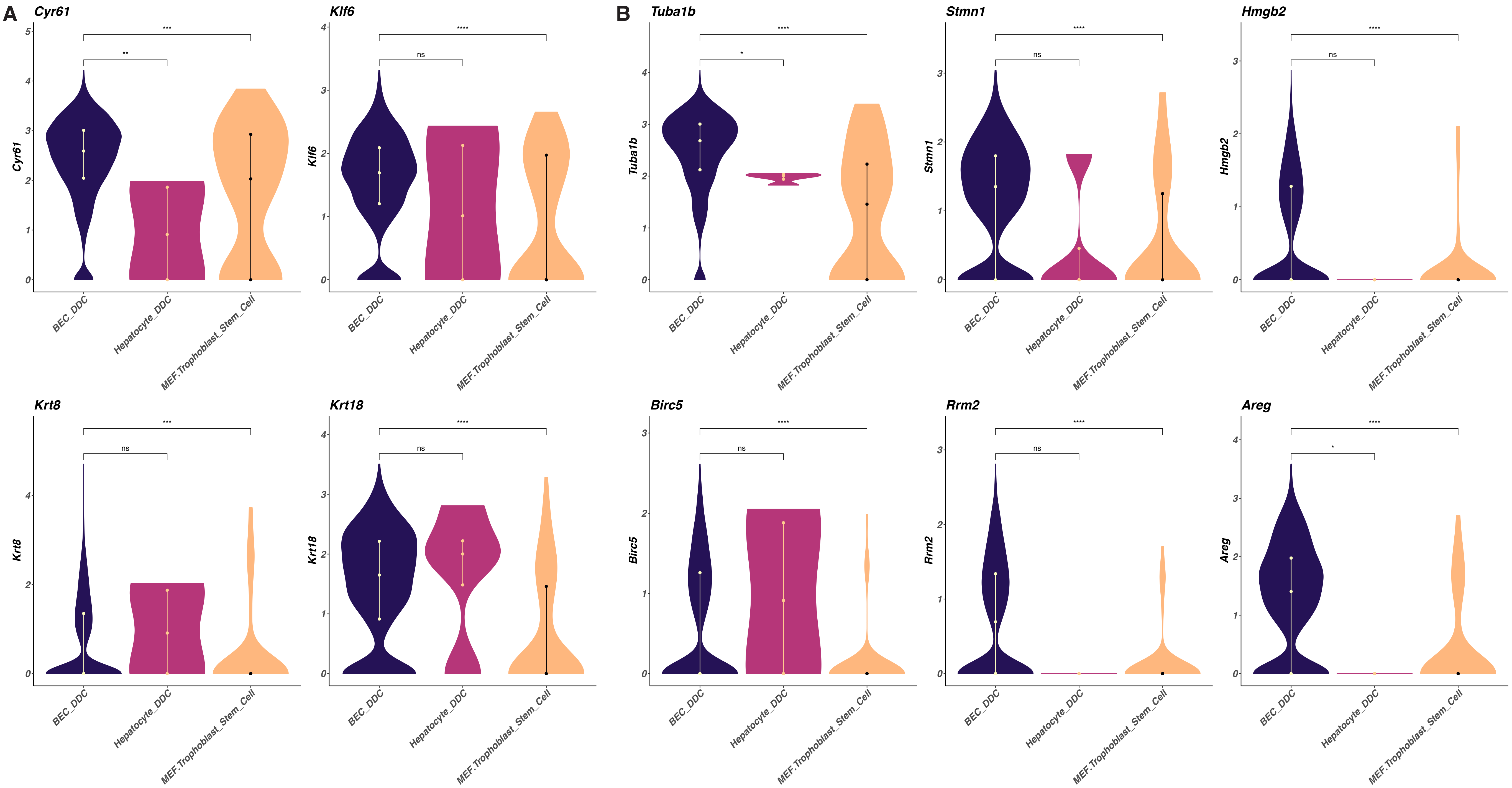
