## Supplementary figures and images for "Capybara: A computational tool to measure cell identity and fate transitions"

### Method Figures

#### Method Figure 1

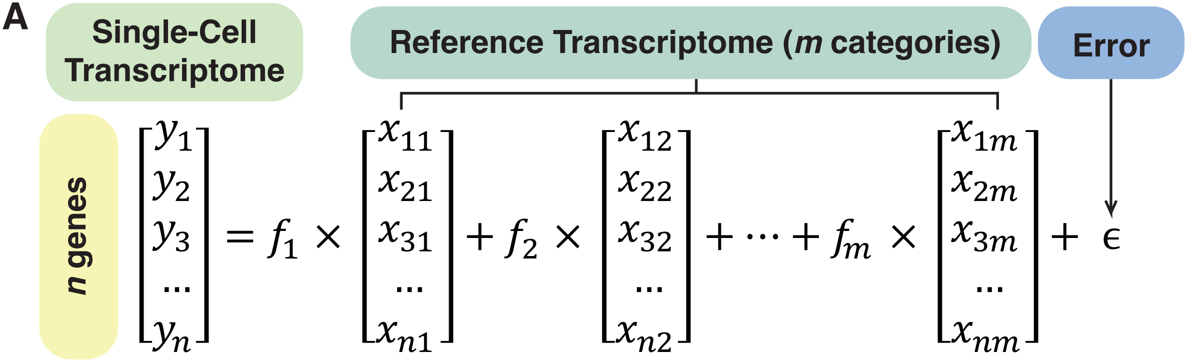

#### Method Figure 2

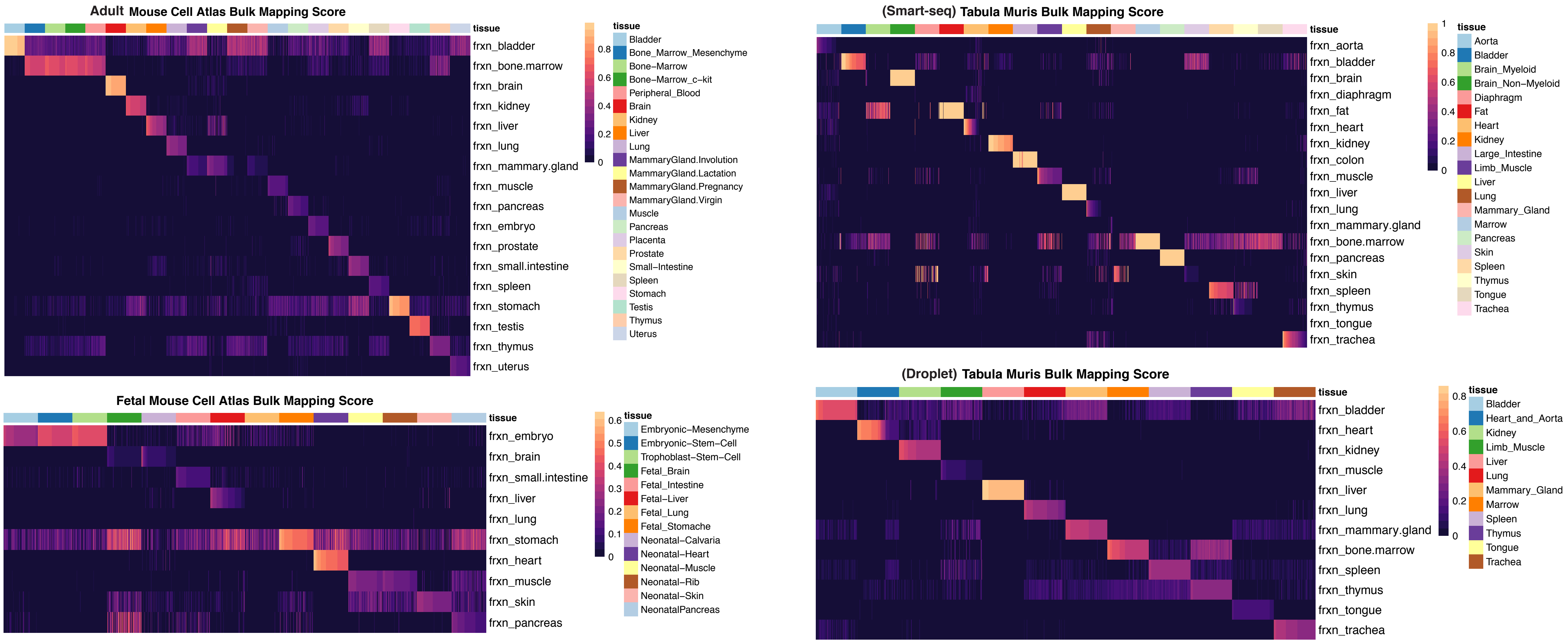

#### Method Figure 3

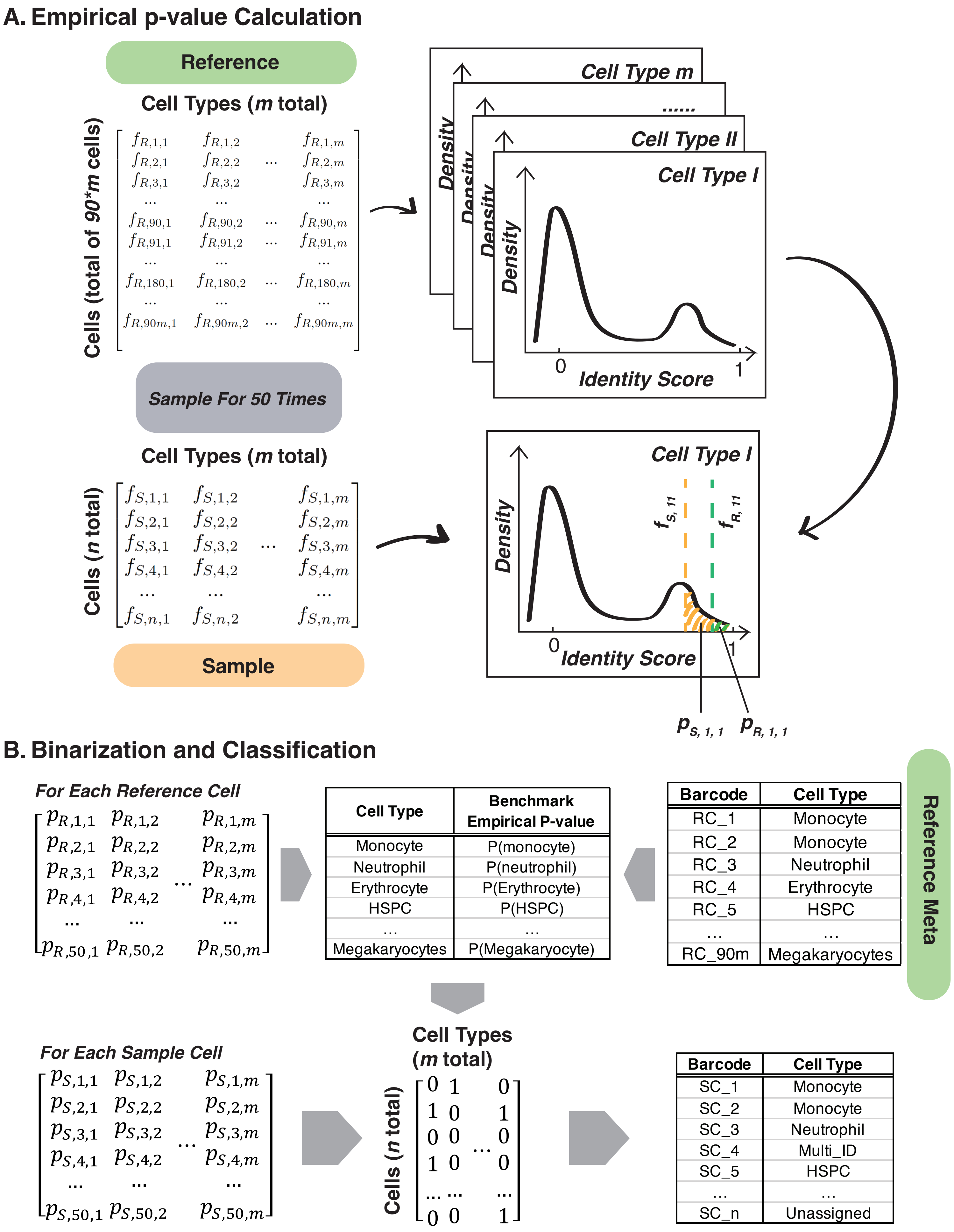
